# Local adaptation of a marine diatom is governed by genome-wide changes in diverse metabolic processes

**DOI:** 10.1101/2023.09.22.559080

**Authors:** Eveline Pinseel, Elizabeth C. Ruck, Teofil Nakov, Per R. Jonsson, Olga Kourtchenko, Anke Kremp, Matthew I.M. Pinder, Wade R. Roberts, Conny Sjöqvist, Mats Töpel, Anna Godhe, Matthew W. Hahn, Andrew J. Alverson

## Abstract

Marine phytoplankton play essential roles in global primary production and biogeochemical cycles. Yet, the evolutionary genetic underpinnings of phytoplankton adaptation to complex marine and coastal environments, where many environmental variables fluctuate and interact, remain unclear. We combined population genomics data with experimental transcriptomics to investigate the genomic basis underlying a natural evolutionary experiment that has played out over the past 8,000 years in one of the world’s largest brackish water bodies: the colonization of the Baltic Sea by the marine diatom *Skeletonema marinoi*. To this end, we used a novel approach for protist population genomics, combining target capture of the entire nuclear genome with pooled sequencing, and showed that the method performs well on both cultures and single cells. Genotype-environment association analyses identified >3,000 genes with signals of selection in response to major environmental gradients in the Baltic Sea, which apart from salinity, include marked differences in temperature and nutrient availability. Locally adapted genes were related to diverse metabolic processes, including signal transduction, cell cycle, DNA methylation, and maintenance of homeostasis. The locally adapted genes showed significant overlap with salinity-responsive genes identified in a laboratory common garden experiment, suggesting the Baltic salinity gradient is a major factor driving local adaptation of *S. marinoi*. Altogether, our data show that local adaptation of phytoplankton to complex coastal environments, which are characterized by a multitude of environmental gradients, is driven by intricate changes in diverse metabolic pathways and functions.

## INTRODUCTION

Given their essential roles in ecosystem functioning (1), understanding how marine phytoplankton adapt to changes in their environment is essential for making predictions on how they will be impacted by environmental change (2). Experimental work showed that phytoplankton species can respond rapidly to environmental change, both through phenotypic plasticity and rapid evolution of metabolic traits (3, 4). However, the genetic underpinnings of marine phytoplankton adaptation to novel and changing environments remain elusive (5), including the evolutionary processes, molecular pathways, and genes that drive adaptive change in these organisms. Unraveling this requires going beyond phenotypic responses to investigate how genotypes change in the face of adaptation. Experimental evolution combined with whole-genome resequencing can provide detailed insights into the rate of adaptation *in vitro*, as well as the genes involved in the early stages of adaptation (6, 7) but experiments are often limited to model strains. Many of these have been grown in culture for decades, which can result in genetic change caused by artificial selection, loss of gene function, and recombination (8–10). Phytoplankton species also exhibit high levels of intraspecific variation in their responses to the environment (11–13), which may not be captured by investigations of model strains. Finally, laboratory treatments cannot fully mimic the complexity of marine ecosystems where many abiotic and biotic parameters fluctuate and interact (14, 15). Studies of natural populations can provide ecological context to laboratory studies of model strains and provide valuable new insights into adaptive change in microeukaryotes (16).

To help fill this knowledge gap, we investigated how a globally distributed marine phytoplankton species, *Skeletonema marinoi*, successfully colonized and adapted to the brackish Baltic Sea (Figs 1a-c). Saline waters from the North Sea inundated the freshwater basin that is now the Baltic Sea some 8,000 years ago, creating the world’s largest inland brackish water body (17). The salinity gradient separating marine and freshwater environments is one of the major ecological divides for microorganisms (18), including diatoms, but the area was nevertheless colonized early on by *S. marinoi* (19). Since its colonization, *S. marinoi* has become an abundant phytoplankton species and prominent primary producer in the Baltic Sea (20). Microsatellite DNA separated *S. marinoi* into high-salinity North Sea and low-salinity Baltic Sea populations, suggestive of reduced gene flow between the two regions (21). Although salinity is generally considered to be the major abiotic factor structuring diversity in the Baltic Sea (22), several other gradients, including temperature and nutrient availability, have likely imposed additional selection pressures on *S. marinoi* in the Baltic Sea.

**Fig. 1.**
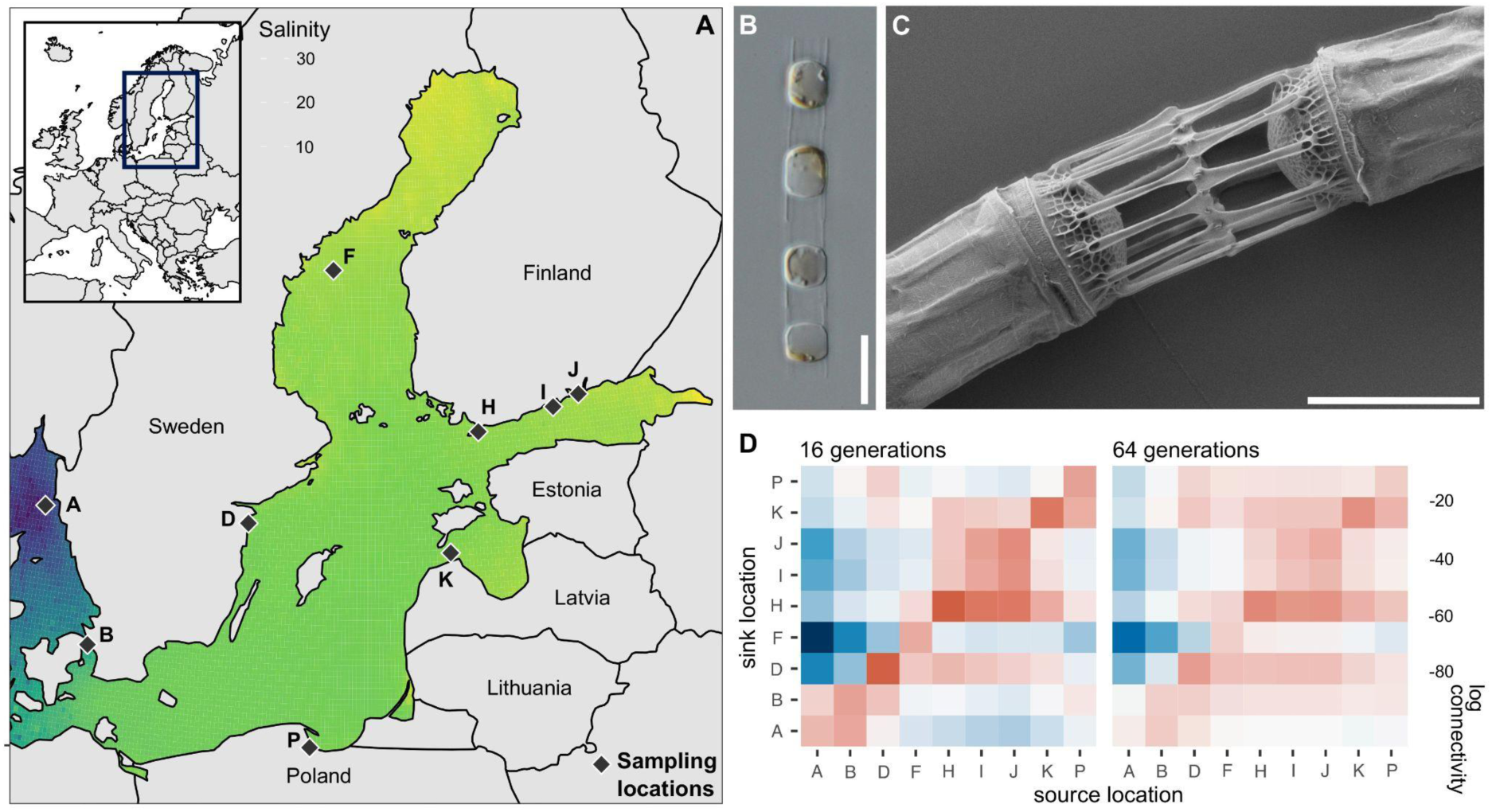
| Sampling locations and study system. **a.** The North Sea - Baltic Sea salinity gradient, with sampling locations for *S. marinoi*. Salinity measurements for the period 2010–2018 used for the interpolation were downloaded from ICES (ICES Ocean Hydrography, 2020. ICES. Copenhagen) and Sharkweb (https://sharkweb.smhi.se/hamta-data/). The inset map in the top left corner shows the broader geographic region. **b.** Light micrograph of a *S. marinoi* culture (scale bar = 10 μm) **c.** Scanning electron micrograph of *S. marinoi* strain RO5AC (scale bar = 5 μm). The micrograph shows the linking spines that connect individual cells, resulting in chain formation. The SEM image was obtained by the Centre for Cellular Imaging at the University of Gothenburg and the National Microscopy Infrastructure, NMI (Sweden, VR-RFI 2016-00968). **d.** Heatmaps showing multigenerational stepping-stone connectivity (16 and 64 generations) between the sampling locations for a particle the size of *S. marinoi*, calculated from the seascape connectivity model. Data were averaged across all months, drift depths, and drift durations. Multigenerational connectivity represents the probability to go from locality X to Y using stepping-stone dispersal over *n* generations. Dark blue values represent lowest connectivity, whereas dark red values represent highest connectivity. Dispersal from locality X to Y is different than going from Y to X, because of possible asymmetric water transport. Connectivity values for the 64-generation heatmap are generally lower than those for the 16-generation heatmap, despite allowing for more generations: in consecutive iterations of the model, particles are lost when dispersing out of the domain (and no new particles are generated), and therefore, multigenerational connectivity needs to be interpreted in a relative sense.

We designed and applied a novel population genomic approach – combining genome capture, single-cell genomics, and pooled sequencing (pool-seq) – to understand how *S. marinoi* locally adapted to the Baltic Sea over the past 8,000 years (Fig. 1a). Our study sheds new light on the evolutionary genetic underpinnings that allow phytoplankton to adapt to complex coastal environments, providing novel and timely insights into the tempo and mode of local adaptation in microeukaryotes (5).

## RESULTS & DISCUSSION

### Leveraging target capture for population genomics

Population genomics on microeukaryotes is challenged by many factors, including large or unknown genome sizes and the limited resolution of traditional methods, such as microsatellites (23). We designed a novel approach to obtain genome-wide nuclear SNPs from hundreds of individuals. Our method combines target capture of the complete nuclear genome, followed by pool-seq to minimize costs. Target capture avoids issues with bacterial contamination and over-sequencing of organellar DNA, redirecting sequencing effort to the target genome. We established *S. marinoi* strains from resting cells in surface sediments collected from the North Sea and Baltic Sea (2010–2018). The genetic composition of these diatom seed banks is locally stable over decades (24, 25), so sample collection over a nine-year period should not significantly impact inferences on population structuring or selection. In total, 245 cultured strains were included in pools from localities B, F, I, J, K, and P, as well as 121 single cells for localities A, D, and H (Fig. 1), where germination success of resting cells was lower (Supplementary Table S1). We achieved read-mapping rates of 96.6–99.0% and recovered 2,197,240 filtered biallelic SNPs across the nine pools (Supplementary Fig. S1). Of these, 1,259,039 SNPs (∼57 %) had a minimum of 20x coverage across all pools. These were used for all subsequent analyses. The high SNP coverage is well-suited for detecting signatures of selection, which stands in stark contrast with the coverage provided by restriction site-associated DNA sequencing (RADSeq) which is usually too low to reliably detect loci under selection (26). Pools assembled from single cells and cultures had comparable SNP coverage (Supplementary Fig. S1), underscoring the utility of our method for uncultivable organisms. Our approach thus performs equally well as a recently designed SAG-RAD protocol that combines single-amplified genomes with RADSeq in microeukaryotes (27), but has the additional advantage of eliminating bacterial contaminants, which can be problematic for the short sequences typical of RADSeq. One drawback of our method is the requirement of a reference genome or transcriptome for bait design. For species with large genomes, such as many dinoflagellates, it will be too costly to capture the entire genome. However, target capture can also be used to sequence specific genes or genomic regions useful for population genomics (28).

### The North Sea and Baltic Sea are home to two distinct populations of *S. marinoi*

Using genome-wide SNP data, we detected two populations. One is confined to the North Sea and Danish Straits (localities A and B: hereon referred to as ‘North Sea’), and the other one is restricted to the Baltic Sea (all other localities) (Figs 2a, c, Supplementary Fig. S2a). As such, the population structure of *S. marinoi* mirrors that of other Baltic organisms, including mussels, fish, and seaweeds (22). We used a biophysical model that accounts for oceanographic currents to estimate the degree of seascape connectivity and predicted dispersal of *S. marinoi* between regions (Fig. 1d). This included the calculation of one- and multi-generation connectivity across 16, 32, and 64 generations and assumed stepping-stone dispersal, which can uncover long-term connectivity between sites (29, 30). Our model highlighted the geographical isolation of the Baltic Sea, as few modeled trajectories of *S. marinoi* can reach the Baltic Sea from the North Sea through surface currents, even when allowing for multi-generational dispersal (Fig. 1d). In contrast, there is high connectivity between localities within each sea (Fig. 1d). We found no significant isolation-by-distance pattern for either the one- or multigenerational connectivities (Mantel test, P-value = 0.10 for the 64-generation model, Fig. 2b). However, there is significant isolation by distance (P-value = 0.004) when using the shortest distance over the sea as a metric for connectivity, but this pattern is driven entirely by the contrast between the North Sea and the Baltic Sea, as there is no significant isolation by distance within the Baltic Sea (P-value = 0.25) (Supplementary Fig. S2b). Together, these analyses indicate that the narrow Danish Straits pose a strong dispersal barrier for *S. marinoi*, as it does for other micro- and macrobiota in the area, including mobile and sessile organisms, as well as drifters (22).

**Fig. 2.**
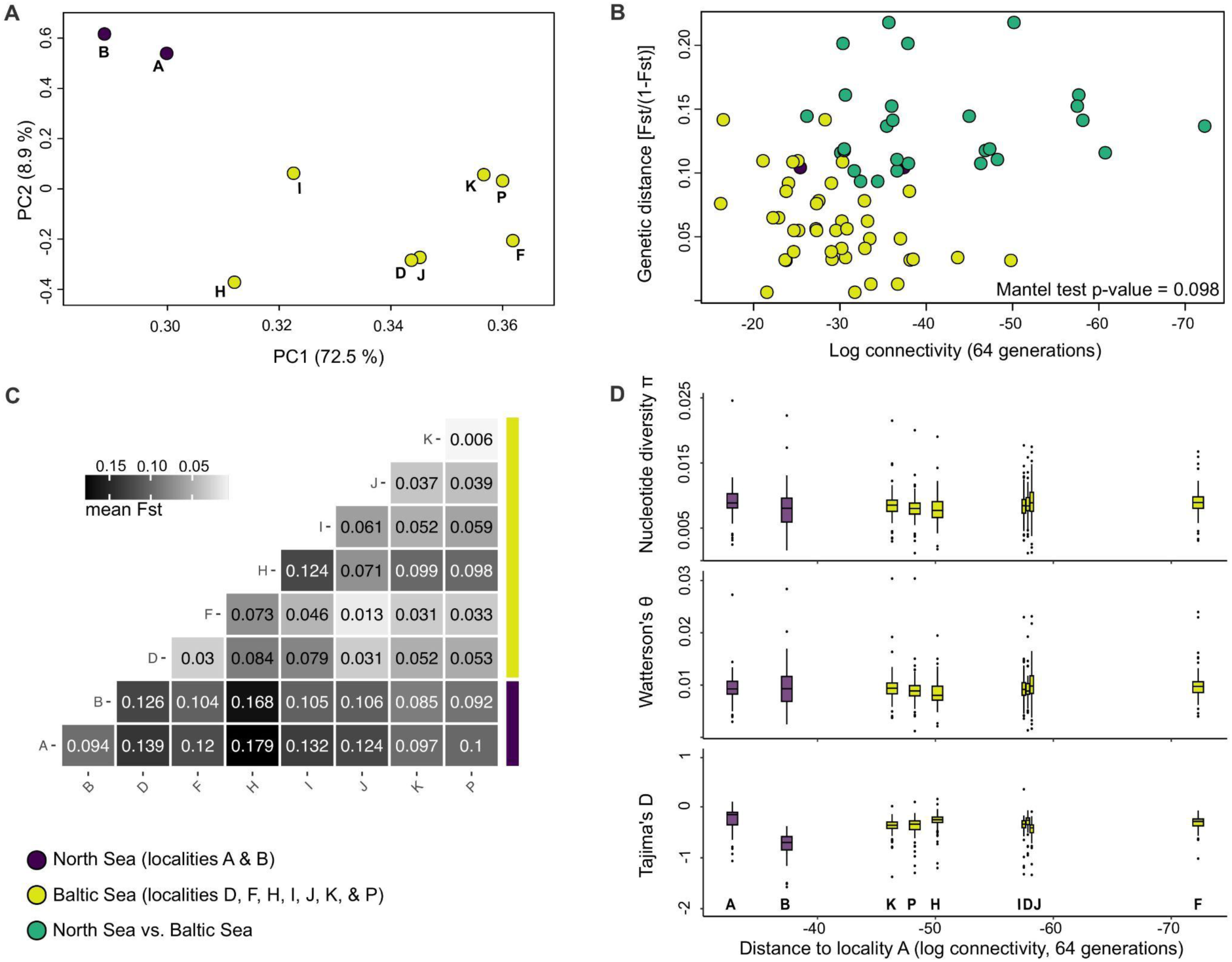
| Population structure of *S. marinoi*. **a.** PCA of the allele frequencies, showing clear distinction between samples from the North Sea and the Baltic Sea. **b.** Isolation-by-distance plot. Distance is measured as the multi-generational stepping-stone connectivity, across 64 generations (see Fig. 1d). Each pair of localities is plotted twice due to asymmetric water transport between localities. **c.** Pairwise genome-wide *F*_ST_ between all localities, showing the lowest levels of population differentiation between localities from the Baltic Sea. **d.** Measures of genetic variation: nucleotide diversity π, population mutation rate θ_W_ (Watterson’s theta), and Tajima’s D. Values were averaged across each contig for each locality, and visualized as boxplots. Outlier SNPs were removed prior to creating the plots in panels a-c.

### Similar levels of genetic diversity in the two populations

Genetic diversity, expressed as nucleotide diversity π and the population-scaled mutation rate *θ*_W_, did not differ between the North Sea and Baltic Sea populations (paired *t*-test; P-value = 0.85 and 0.22, respectively), indicating similar levels of genetic diversity in the two regions (Fig. 2d). Although Tajima’s *D* was significantly lower in the North Sea (P-value < 1e-5), this pattern was driven by locality B (Fig. 2d). Clearly, despite population differentiation and its relatively confined setting, the Baltic Sea population does not exhibit reduced genetic diversity. This suggests that if *S. marinoi* experienced a population bottleneck when colonizing the Baltic Sea, the bottleneck was either too small, or too long ago, to be detected today. Although microeukaryotes have short generation times and are assumed to harbor extraordinarily high levels of intraspecific variation (31), the genetic diversity of *S. marinoi* in our study area is comparable to populations of small mammals, insects, and plants (32). Altogether, our data indicate that Baltic *S. marinoi* is not a sink population, as would be expected from a maladapted population sustained only through constant migration from a diverse source population. On the contrary, it is likely well adapted to the Baltic Sea.

### Signatures of local adaptation to the Baltic Sea

We used genotype-environment association (GEA) to detect outlier SNPs correlated with seasonal environmental variables in the entire study area. However, the environmental differences between the North and Baltic seas are correlated with *S. marinoi* population structure (Supplementary Fig. S3). This challenges the ability of GEA approaches to control for neutral patterns of population structuring (33, 34), so we restricted GEA analyses to the Baltic Sea. To avoid issues with collinearity between environmental variables, we used the first two principal components of a Principal Component Analysis (PCA) on the Baltic environmental variables, which explained 72% of the variation, as input for a latent factor mixed model, LFMM (35) and Bayesian approach, BayPass (36) (Supplementary Figure S4). PC1 and PC2 correlated with the north-south and east-west environmental gradients of the Baltic Sea, respectively. Although the environmental dataset was not exhaustive, our GEA analysis indirectly accounts for unmeasured variables. Sea-ice variation and composition, for instance, follows a north-south gradient in the Baltic Sea (37).

We found that most outlier SNPs were associated with the north-south gradient (LFMM: 9,819, BayPass: 396), suggesting that differences in salinity, summer temperature, and nutrient availability impose the greatest selective pressures in the area. The east-west gradient (LFMM: 155, BayPass: 771 outlier SNPs) was correlated with differences in salinity, temperature (winter, spring, autumn), pH, alkalinity, light availability, oxygen, nitrate, and silicate. Outlier SNPs were distributed across the genome (Figs 3d, Supplementary Figure S5c). Combining LFMM and BayPass results, we recovered 10,019 outlier SNPs for the north-south gradient and 900 for the east-west gradient, representing 3,747 and 555 genes, respectively (Fig. 3a). Most GEA-outliers were located in exons, and about half of these were missense SNPs (Fig. 3b-c). These outliers are the most likely to be under selection or linked with SNPs that are under selection. Despite the challenges of distinguishing the two, here and elsewhere in our study we focus on SNPs associated with changes in protein sequence (e.g., missense SNPs) as targets of selection. We also explored SNPs in untranslated regions (UTRs), as these regions affect the translation, degradation, and localization of mRNAs (38).

**Fig. 3.**
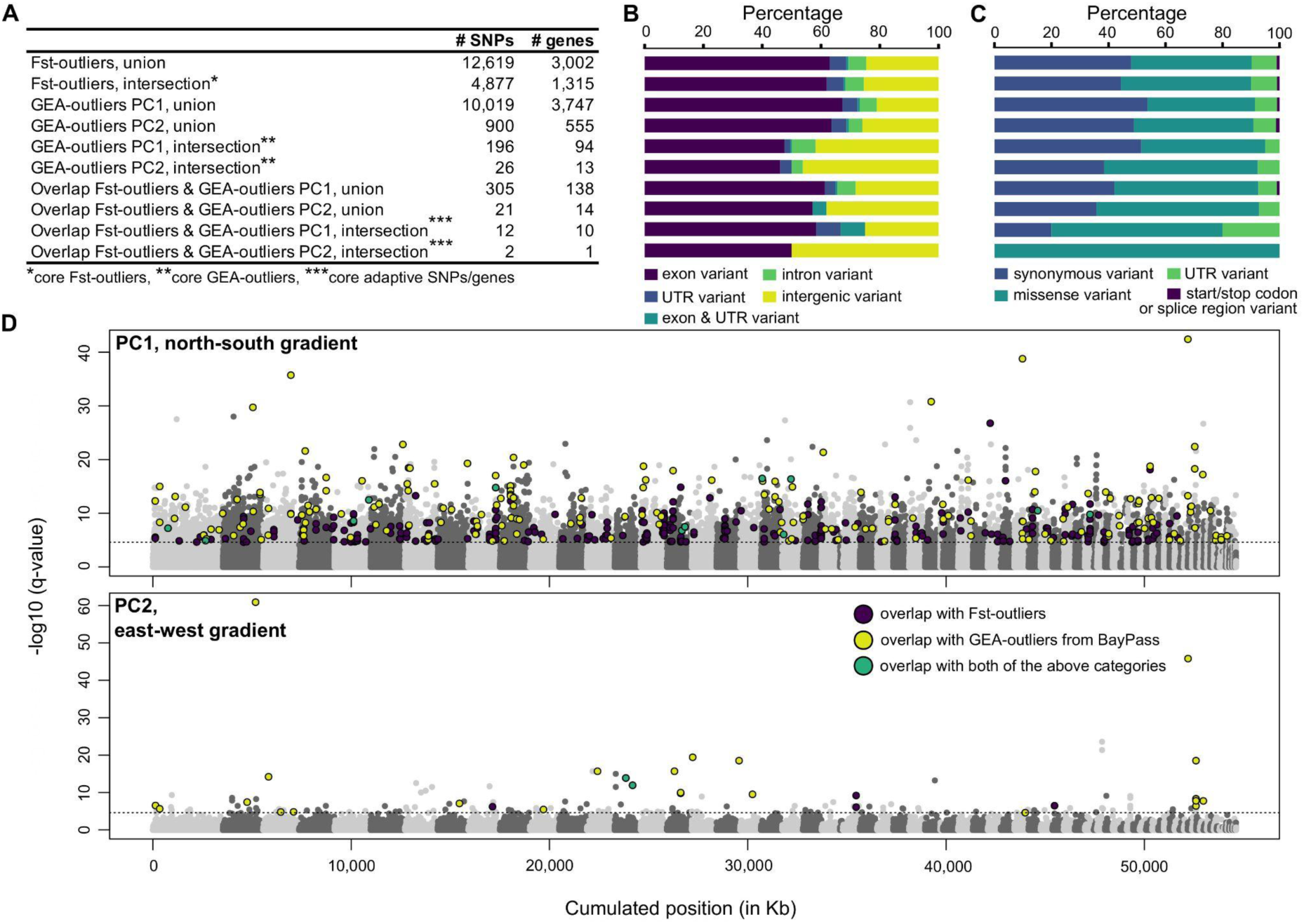
| Outlier SNPs and genes associated with the Baltic Sea environmental gradients. **a.** Table showing the number (#) of outlier SNPs and genes in each tested category. *Union* refers to the full set of SNPs or genes found by one or both approaches (i.e., BayPass/FET for the *F*_ST_-outliers or LFMM/BayPass for the GEA-outliers). Similarly, *intersection* refers to outlier SNPs and genes that were part of the overlap of both approaches in each category. The bottom four rows refer to outlier SNPs or genes that overlapped between the union and intersection lists of the *F*_ST_- and GEA-outliers, listing both PC axes separately. **b.** Types of outlier SNPs for the categories in (**a**). The percentage was calculated using the full set of outlier SNPs. The labels of the vertical axis correspond with the labels in (**a**). **c.** Types of outlier SNPs for the categories in (**a**), only showing SNPs located in exons. The percentage was calculated using the total number of outlier SNPs located in exons. The labels of the vertical axis correspond with the labels in (**a**). **d.** Manhattan plots showing outlier SNPs associated with the environment of the Baltic Sea as estimated by LFMM, shown separately for both PC axes. The dotted line represents the 1% FDR significance threshold. Different contigs are indicated with alternating shades of gray. The colored dots represent outlier SNPs that overlap with *F*_ST_-outliers (dark blue), BayPass’ GEA test (yellow), or both (green).

Genes with outlier SNPs, and thus possible signals of local adaptation, had functions that were clearly associated with the Baltic environmental gradients. For instance, we found selection on genes involved in cation, sodium, and phosphate transmembrane transport, chemical homeostasis, and oxidative stress along the north-south gradient (Fig. 4a, Supplementary Fig. 6a, c-d), indicative of changes in how the diatom responds to nutrient availability and osmotic (salinity) stress. In addition, histone monoubiquitination was significantly enriched along the north-south gradient (Supplementary Fig. 6d), suggesting a possible role of epigenetics in local adaptation, similar to vascular plants (39, 40). Outlier genes along the east-west gradient were involved in energy homeostasis and response to abiotic stimuli, including light intensity (Fig. 4b, Supplementary Fig. 6b). The east-west gradient is associated with Secchi depth, a measure of water turbidity, which suggests that light availability could drive adaptive change. However, here and elsewhere, it is possible that covariate stressors evoke similar responses. In addition, we found significant enrichment of proteolysis, macroautophagy, and organelle disassembly along the east-west gradient (Fig. 4b, Supplementary Fig. 6b), possibly underscoring selection on intracellular breakdown and recycling of cytoplasmic compounds and organelles, similar to the acclimated response to low salinity in *S. marinoi* (11). The east-west gradient was further associated with local adaptation on genes involved in the electron transport chain, possibly reflecting changes in energy metabolism (Fig. 4b).

**Fig. 4.**
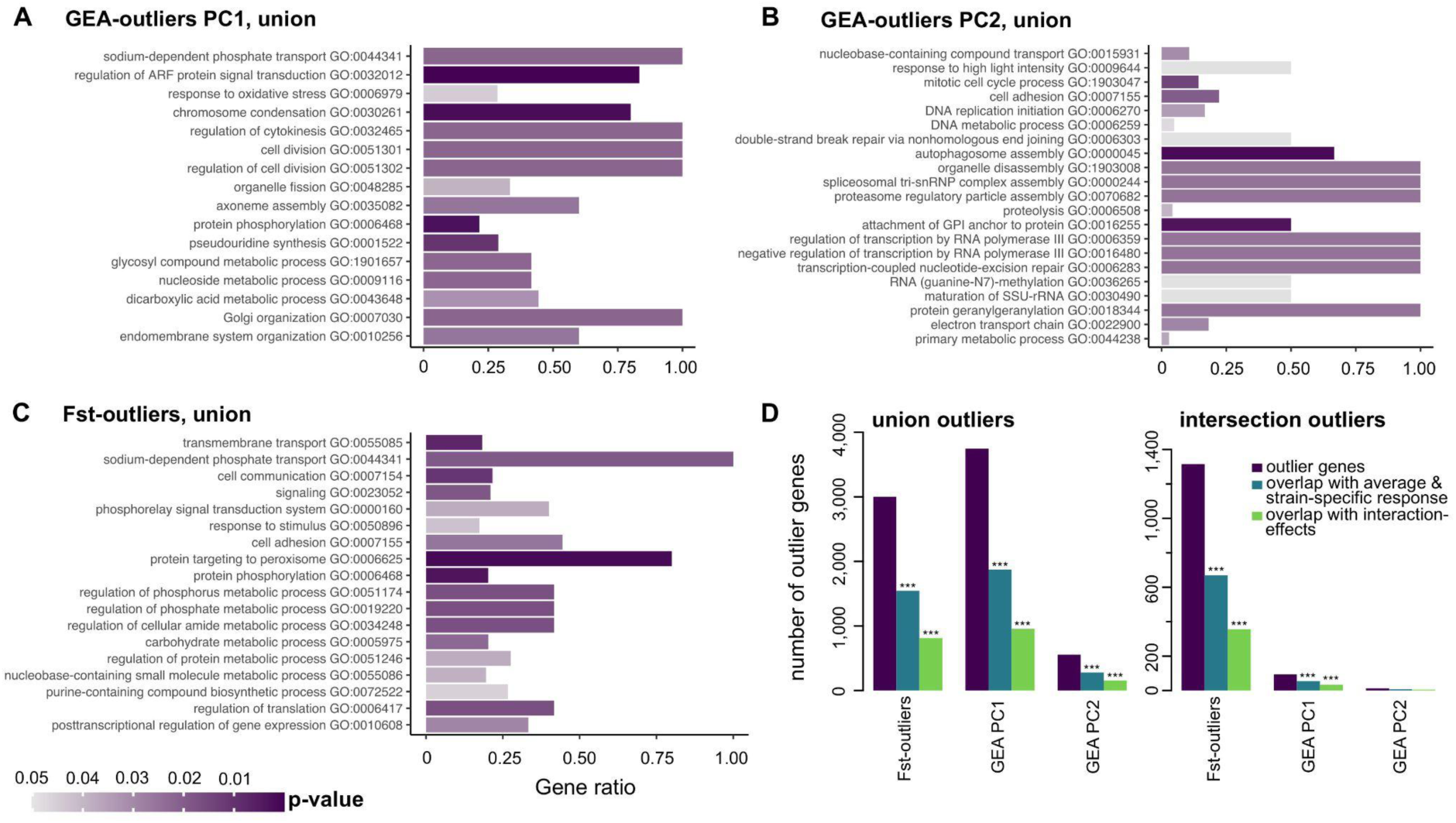
**| a-d.** Barplots showing GO enrichment results for the GEA tests for PC1 (north-south gradient) (**a**) and PC2 (east-west gradient) (**b**), showing the union between both approaches (LFMM/BayPass), and *F*_ST_-outlier genes (**c**), showing the union between both approaches (BayPass/FET). GO enrichment was run on genes with at least one missense outlier SNP. Bars are colored by topGO’s P-value. For each plot, GO terms include Biological Process GO terms that were retained after running REVIGO. The height of the bars indicates the proportion of genes with a given GO term that are enriched relative to the total number of genes with this GO term in the genome of *S. marinoi*. **d.** Barplots showing the overlap between the sets of outlier genes from this study and differentially expressed genes in *S. marinoi* in response to low salinity. Outlier genes are subdivided in different categories: *F*_ST_-outliers, GEA-outliers for PC1, and GEA-outliers for PC2. The *union* barplot indicates the full set of outlier SNPs detected by one or both approaches for each category, whereas the *intersection* barplot indicates the intersection between the outlier tests. Gene expression data were obtained from eight strains, originating from localities A, B, D, F, I, J, K, and P, which were exposed to salinities mimicking the Baltic Sea salinity cline (24, 16, and 8). We tested for differentially expressed (DE) genes for each salinity contrast (8-16, 16-24, and 8-24) within each strain and by combining data of all strains, resulting in a total of 7,676 differentially expressed genes (= *average & strain-specific response* in the plot). For all combinations of strains, we also tested for interaction-effects for each salinity contrast, thus testing for significant strain-specific responses: this resulted in 3,958 differentially expressed genes (= *interaction-effects* in the plot). Significant overlap between the outlier and differentially expressed genes are indicated with asterisks (*** = P-value < 0.005).

Diatoms undergo progressive cell size reduction by mitosis until the cell size reaches a species-specific size threshold which, often together with an environmental trigger, prompts sexual reproduction to restore maximal cell size (41). In *S. marinoi*, shifts to lower salinities can trigger sexual reproduction

(42). Similarly, unfavorable conditions induce formation of resting cells that can survive in bottom sediments for more than a century (25). Although many of the genes involved in these processes are unknown, they likely involve cell cycle and cell division genes, which were under selection along both the north-south and east-west gradients (Fig. 4a-b, Supplementary Fig. 6a-d). In addition, we found several meiotic genes under selection (e.g., meiotic recombination protein SPO11-2). This suggests that local adaptation to the Baltic Sea involves selection on growth rates, resting cell formation, and/or frequency of sexual reproduction. This is consistent with laboratory experiments that showed that Baltic strains evolved higher growth rates relative to North Sea strains through countergradient selection to cope with the demanding Baltic Sea environment, but these higher growth rates come at a cost when competing in ancestral waters (43).

The overlap between outlier SNPs detected by LFMM and BayPass is significantly larger than expected by chance (hypergeometric test, P-value < 0.0001) with 196 shared SNPs (94 genes) for the north-south gradient, and 26 shared SNPs (13 genes) for the east-west gradient (Fig. 3a, Supplementary Fig. S5a). These genes carry the strongest evidence for association with the Baltic gradients and were involved in diverse cellular functions, including signal transduction, fatty acid biosynthesis, the cell cycle, energy homeostasis, and ion channels. Most notably, several of these core outliers belonged to genes involved in vitamin B6 (north-south gradient) and glutathione metabolism (east-west gradient), both of which are involved in defense mechanisms towards external stressors, including reactive oxygen species (44). We also detected a chitin synthase and a protein with chitin-binding activity (north-south gradient). Chitin is thought to be important for low salinity tolerance by affecting cell wall remodeling and/or buoyancy adjustments, as was suggested for the euryhaline diatom *Cyclotella cryptica* during acute hyposalinity stress (45). In addition, chitin might play a role in cell linkage and chain formation in *S. marinoi* (46). Given the importance of chain length to grazer susceptibility (47), selection on chitin genes suggests that variation in predation (48) might also invoke variable selection pressures across the Baltic Sea (49).

Local adaptation of *S. marinoi* to the Baltic Sea gradients is also likely associated with population structure at fine geographic scales. Specifically, two Baltic localities (H and I) showed relatively high *F*_ST_-differentiation from the other Baltic sites (Figs 2a, c), despite high oceanographic connectivity (Fig. 1d). Microsatellite studies on Baltic *S. marinoi* also showed pronounced genetic differentiation along a south-west to north-east trajectory during a single bloom-period (20), and revealed significant genetic differentiation over only tens of kilometers (50). The mechanism by which this happens could involve priority effects, which have been found to be important in microcosm experiments (51, 52). Priority effects entail a scenario of rapid population growth after initial colonization, which saturates the open niche, and is followed by rapid local adaptation (50–52). The local sediment seed banks of *S. marinoi* are thought to further promote local adaptation by increasing the standing genetic variation (53), and sustain locally adapted variants through time, buffering against later immigrants and maintaining population differentiation (50–52).

### What enabled *S. marinoi* to colonize the Baltic Sea?

We found a large number of SNPs with significantly different allele frequencies in the North versus Baltic Seas by applying the BayPass model and Fisher’s Exact Test (FET) (hereon referred to as ‘*F*_ST_-outlier approaches’). Taken together, BayPass and FET detected 12,619 SNPs in 3,002 genes. The overlap between both approaches was significant (hypergeometric test, P-value < 1e-5) and included 4,877 SNPs (1,315 genes, Fig 3a). Similar to the GEA tests, SNPs were distributed across the genome and were mostly located in exons, with about half representing missense SNPs (Fig. 3b-c, Supplementary Fig. S7). The small number of SNPs that were both *F*_ST_- and GEA-outliers (Fig. 3a) are strong candidates for a role in the range expansion of *S. marinoi* in the Baltic Sea, as they show large allele frequency differences between the regions and are also associated with the Baltic environmental gradients. These included a chitinase and genes involved in, amongst others, signal transduction, cell cycle, fatty acid and glutathione metabolism, oxidative phosphorylation, and stress defense. A gene encoding histone arginine demethylase contained multiple outlier SNPs, which suggests that evolution of DNA methylation played a role in range expansion of *S. marinoi*.

We also explored the larger set of *F*_ST_-outliers, regardless of overlap with GEA, to gain a general idea of genes and pathways that might have enabled colonization of the Baltic Sea (Fig. 4c, Supplementary Fig. S6e-j). Most strikingly, these *F*_ST_ outliers are enriched for signal transduction, cell communication, response to external stimuli, and defense response. This includes various protein kinases: (i) calcium/calmodulin-dependent protein kinases which are differentially expressed in *S. marinoi* in response to hyposalinity (11) and which might play a role in osmotic sensing in diatoms (54), (ii) serine/threonine protein kinases, including *mTOR*, which have been involved in stress responses and resting cell formation in diatoms (55, 56), (iii) cGMP-dependent protein kinases, which play a role in the salt stress response of plants (57), and are differentially expressed in diatoms in response to copper exposure and deficiency (58), (iv) histidine kinases, which are involved in signal transduction across the cell membrane, nutrient sensing, light perception, and oxidative and osmotic stress response in plants, fungi, and algae (59), and lastly, (v) cAMP-dependent protein kinases which rely on the ancient signaling molecule cAMP that serves as a stress indicator across the tree of life (60). Several of the histidine kinases were also GEA-outliers. Clearly, widespread changes in signal transduction pathways that are involved in various metabolic processes, including the stress response and osmotic regulation, could have played a central role in *S. marinoi*’s range expansion.

The Baltic Sea is naturally vulnerable to nutrient enrichment due to stratification and long retention times (61). As a result, during its initial colonization, *S. marinoi* may have been confronted with both lower salinities and higher nutrient concentrations. Furthermore, during the last century, the Baltic Sea has undergone large-scale eutrophication due to increased anthropogenic inputs (61), and it is conceivable that these changes add additional selection pressures on Baltic *S. marinoi* today. Several genes associated with nitrogen and phosphorus utilization were enriched in the *F*_ST_-outliers (Fig. 4c, Supplementary Fig. S6e-j). *F*_ST_-outliers were further enriched for processes related to carbohydrate, lipid, and polyamine metabolism (Fig. 4c, Supplementary Fig. S6e-j), all of which are differentially expressed in response to hyposalinity in *S. marinoi* (11). Altogether, our data indicate that adaptation of *S. marinoi* to the Baltic Sea balanced an array of selection pressures, involving differences in osmotic pressure, nutrient availability, and the general impact of a suboptimal, potentially stressful, environment.

### What is the role of salinity in adaptation to the Baltic Sea?

The salinity gradient is one of the most pronounced environmental gradients that structures biodiversity in our study area (22). However, given that salinity is correlated with numerous environmental variables, it is challenging to disentangle the specific role of salinity in driving adaptation and population differentiation. To overcome this issue, we combined our SNP dataset with a common garden experiment that we previously performed. In this experiment, we exposed eight *S. marinoi* strains originating from the same sampling localities as this study to salinity treatments that mimicked the Baltic Sea salinity cline (24, 16, and 8 ppt) (11). For most outlier categories, there was significant overlap between the outlier genes identified in this study and ones that were differentially expressed in response to salinity (hypergeometric test, P-value < 0.005) (Fig. 4d). However, we did not find a relationship between the strength of gene expression differences and the magnitude of between-locality allele frequency differences of SNPs within these shared genes (Supplementary Fig. S8). Thus, genes with outlier SNPs that showed the largest allele frequency differences between localities did not consistently show the strongest differences in gene expression between *S. marinoi* strains that originated from the same localities.

Among the set of 33 differentially expressed genes considered essential to low salinity acclimation in *S. marinoi* (11), 16 were recovered as outlier genes, of which 11 had missense and/or UTR SNPs. These included genes involved in fatty acid and polyamine metabolism, the glycine/serine/threonine pathway, and transporters for solutes, amino acids, polyamines, and cations.

The full set of differentially expressed genes identified by Pinseel et al. (11) that also contained outlier SNPs (missense/UTR) were involved in several other processes that are important for salinity acclimation, including oxidative stress responses, polyamine biosynthesis, heat shock proteins and calmodulins, chlorophyll biosynthesis, the non-mevalonate pathway, and genes involved in the metabolism of amino acids, including the osmolyte taurine (Supplementary Fig. S9). Although several of these pathways could be important for responses to other stressors, our data suggest that salinity is likely to be a major, though not sole, driver of local adaptation to the Baltic Sea. Finally, a phylogenomic study of Thalassiosirales, the diatom lineage to which *S. marinoi* belongs, identified 532 hemiplasious genes, i.e., ones with persistent ancestral polymorphisms associated with marine–freshwater transitions (62). A total of 151 of our outlier genes were also present in this set of hemiplasious genes, suggesting that the genes which contributed to ancient marine–freshwater transitions may have been similarly important for salinity adaptation over the microevolutionary timescales of this study.

## CONCLUSION

The adaptive potential of phytoplankton species is routinely studied by measuring changes in phenotypic traits in laboratory experiments. We investigated the genetic basis of adaptation by a marine planktonic diatom to its natural environment using a combination of high-resolution population genomics and experimental transcriptomics (11). Our data show that the complex environment of the Baltic Sea, which is characterized by multiple, interacting environmental gradients, drives intricate genome-wide changes in diverse pathways in *S. marinoi*. Given the existence of seasonal variation in most abiotic parameters, it is likely that selection is mostly fluctuating in nature, resulting in multiple coexisting genotypes adapted to different environmental conditions (20). Although transcriptome data indicated that the marine-brackish salinity gradient is a major driver of adaptive change, it is clear from the patterns of selection across the genome that other factors, including nutrient availability, are also important. Thus, our data indicate that the adaptive potential of marine planktonic diatoms hinges at least partially on the ability to rapidly adapt through complex polygenic changes, combined with plastic and evolved changes in gene expression (11). Altogether, this study represents a major step forward in our understanding of the genetic basis of natural adaptive processes in marine phytoplankton, and indicates that if we are to uncover the full adaptive potential of phytoplankton populations we need to account for the polygenic nature of adaptation to complex environments. Ultimately, in-depth insights into the genetic drivers behind microeukaryote adaptation in natural environments will contribute to understanding and predicting their responses to environmental change and vulnerability to extinction (63). The latter is especially important for marginal populations in areas that are forecasted to experience massive environmental change in the coming decades (64, 65).

## MATERIALS AND METHODS

### Sample collection and culturing

Between 2010 and 2018, we collected surface sediment samples from nine localities spanning the Baltic Sea salinity cline (Fig. 1a) and stored these in the dark at 5 °C. This included two localities in the North Sea and Danish Straits, and seven localities in the Baltic Sea. A plankton sample was also obtained for location P (Fig. 1a). We germinated resting cells of *S. marinoi* into monoclonal cultures that were kept at their native salinity, 12 °C, and a 12:12h light:dark light regime (30 μmol photons m_-2_ s_-1_ light intensity). Strains that were kept at salinity 5 or 8 were grown in WC medium (73) with added salts, whereas strains maintained at salinity 16 or 24 were grown in L1 medium (74) with adjusted salinity (11).

### Genotyping *S. marinoi* strains

The taxonomic identity of each strain was confirmed by sequencing the LSU rRNA gene (D1-D2 region), thus ensuring all isolates belonged to *S. marinoi*. LSU was chosen because differentation in this gene corresponds with species boundaries in the genus *Skeletonema* (75). To this end, we harvested cells by centrifugation and flash freezing in liquid nitrogen after which they were stored at -80 °C until DNA extraction. Frozen cells were broken with 1.0 mm glass beads (Biospec Products, OK, USA) by shaking the tubes for 30 seconds at 3,000 OSM in a Minibeadbeater (Biospec Products, OK, USA). We extracted DNA with the DNeasy Plant Mini Kit (Qiagen, Hilden, Germany), after which the 28S region was amplified following the protocol outlined in (11). Single-cell sequencing was necessary for some uncultivable strains. For these, we first performed a whole-genome amplification step using Qiagen’s REPLI-G MiniKit before the 28S amplification. Forward strands were sequenced at Eurofins Genomics (Louisville, KY, USA) and analyzed with Sequencher v5.1 (Gene Codes Corporation, Ann Arbor, MI, USA). Strains were only included in the Pool-seq if their D1-D2 LSU rRNA sequence (567 bp) was 100% identical to the LSU sequence of the *S. marinoi* reference genome (strain RO5AC). Consequently, all *S. marinoi* strains in this study had identical 28S D1-D2 sequences.

### Pool-seq DNA extraction and sequencing

Prior to DNA extraction, cultures were filtered and concentrated in a pellet, then flash frozen in liquid nitrogen. For each strain, ∼40 million cells were harvested, with the exception of strains from locality J where ∼25 million cells were harvested. Cell concentrations were determined using a Benchtop B3 series FlowCAM cytometer (Fluid Imaging Technologies), and an in-house machine learning tool (available on Zenodo) that estimates the number of cells per chain from image data. DNA was extracted with a Qiagen DNeasy kit, DNA quantity and quality were checked with a Qubit 2.0 (Invitrogen) and a TapeStation 2200 (Agilent Technologies), after which DNA of all strains per locality was pooled at equimolar concentrations. In total, DNA of 18 to 41 strains per locality was pooled (Supplementary Table S1). For some localities, cultures grew poorly, resulting in insufficient biomass for standard DNA extraction (Supplementary Table S1). For these localities, we performed single-cell isolations, followed by a whole-genome amplification step with Qiagen’s REPLI-G MiniKit (see previous section), following the manufacturer’s protocol for blood and cells. The DNA quality was assessed as described above then pooled per locality at equimolar concentrations. In total, DNA of 40 to 41 single cells was pooled per locality (Supplementary Table S1).

After DNA pooling, we performed target-capture using a custom-designed probe kit with a SeqCap EZ Workflow (Roche, Indianapolis, IN, USA), which was designed to capture the entire nuclear genome of *S. marinoi*. The capture set was developed using the reference genome of *S. marinoi* strain RO5AC v1.1 (11). For the target-capture, we followed the manufacturer’s protocol, but with half the reaction volume, using a KAPA HyperPlus kit for library preparation, followed by hybridization of the pre-capture library to a SeqCap EZ Probe Pool (Roche). Following target capture, libraries of pooled individuals were sequenced (whole-genome shotgun sequencing, Pool-seq) on an Illumina HiSeq 4000 at the University of Chicago Genomics Facility (paired-end, 100 bp). This resulted in ∼170–260 million reads per pool.

### Raw read processing and SNP calling

Raw sequence reads were trimmed and filtered using Atria v3.1.0 (76), removing adapter sequences and reads shorter than 36 base pairs, and using a minimum quality score of 20 and a quality kmer score of 5. Processed reads were mapped to the *S. marinoi* strain RO5AC reference genome v1.1.2 (available from our Zenodo repository) with BWA-MEM v0.7.17 (66). Note that upon finalizing this study the final assembly, v1.1.8, of *S. marinoi* strain RO5AC became available (GenBank: JATAAI010000000). Ambiguously mapped reads (mapping quality < 20) were removed using SAMtools v1.10 (77). We subsequently used Picard v2.26.10 (http://broadinstitute.github.io/picard/) to sort the reads in coordinate-order and remove PCR duplicated reads. After mapping and quality steps, we retained roughly 44–104 million reads per pool. Preliminary analyses showed no differentiation between the sediment and plankton samples from locality P so they were combined. Next, we used Picard to add read group tags and performed indel realignment in GATK v3.5 (78). A mpileup of the nine individual pools was created with SAMtools mpileup using a minimum base quality of 20, after which indel regions were identified and removed with the *identify-indel-regions.pl* and *filter-sync-by-gtf.pl* scripts from PoPoolation2 (67) using an indel window of 5 and a minimum count of 1. SNP calling was done with PoolSNP (68), downloaded on October 5th 2022, with support of GNU parallel (79). We evaluated three different filtering strategies, using different settings for the minimum coverage (MIN-COV) and minimum allele count (MAC): (i) liberal (MIN-COV 10, MAC 3), (ii) intermediate (MIN-COV 20, MAC 3), and (iii) conservative (MIN-COV 20, MAC 4). In all cases, minimum allele frequency equaled 0.001, and maximum coverage equaled twice the average sequencing depth, individually defined per contig per pool. We used the *filter-pool-seq-sync.R* script, modified from PoolParty’s *r_frequency.R* script (80), to further remove multiallelic SNPs and calculate allele frequencies, all using R v4.1.0. Principal Component Analysis (PCA), Neighbor-joining (NJ) trees, and population differentiation, measured as the fixation index *F*_ST_ (see below for details), showed virtually identical patterns across the three filtering strategies. To minimize false positive detection of outlier SNPs, we restricted all further analyses to the conservative filtering strategy.

### Simulation of seascape connectivity and dispersal

The seascape connectivity model simulates dispersal trajectories with a Lagrangian particle-tracking model, TRACMASS (81), driven off-line with flow velocity fields (3h temporal resolution) from the ocean circulation model NEMO-Nordic (82). The model grid has a horizontal spatial resolution of 3.7 km (2 NM) with 84 vertical levels, a free surface allowing the grid boxes to stretch and shrink vertically to allow for tides, and the atmospheric forcing is based on reanalysis of the ERA40 dataset (83). At the model boundary, tidal harmonics define the sea surface height, and Levitus climatology defines temperature and salinity (84). Climatological data from a number of different databases for the Baltic Sea and the North Sea provided freshwater runoff. Validation of the NEMO-Nordic model has shown that the biophysical model is able to correctly represent the sea surface height, both tidally induced and wind driven (82).

The biophysics model explored the effects of different traits on single-generation dispersal and multigenerational stepping-stone connectivity to identify possible barriers to gene flow in the study area (21, 85). Specifically, the particle-tracking model was parameterized with two drift depths (surface at 0-2 m, and a deeper layer at 10-12 m), two drift durations (10 and 20 days), and with release of virtual particles once every month, modeling transport of suspended *S. marinoi*. Model results are averages over 8 years (1995–2002) representing the range of the North Atlantic oscillation cycle (86) that correlates well with large-scale circulation patterns in this region. Particles were released from all 34,036 model grid cells with a water depth ≤ 100 m, with a total number of 6.4 x 10^8^ released particles. Dispersal probability between all 34,036 grid cells were calculated and summarized in a connectivity matrix (87). From the connectivity matrix we calculated the multi-generation connectivity assuming stepping-stone dispersal within the model domain. Multigeneration connectivity was calculated by multiplication of the connectivity matrix across 16, 32, and 64 generations, expanding upon the number of generations of previous work which observed that the largest number of generations tested (32) showed the highest correlation between oceanographic trajectories and directional relative migration assessed from microsatellite data (21). The resulting connectivity matrices were summarized and visualized in R. To summarize the results, we averaged the data across the two drift depths and durations as well as all months for each generation (see Fig. 1d for the plots for the 16- and 64-generation models: the full set of plots can be found on Zenodo).

### Population structure and genetic variation

We investigated population genetic structure using a subset of SNPs that were not considered outliers (identified using the approaches in the following two sections). As such, we removed 23,177 outlier SNPs from the dataset prior to calculating population structure. An NJ tree and PCA were constructed using PoolParty’s *r_structure.R* script, using a window size of 1,000, and 1,000 bootstraps for the NJ tree. The *F*_ST_ of putatively neutral SNPs was calculated in Poolfstat (69). We calculated both pairwise *F*_ST_ (between each combination of sampling sites) and sliding-window *F*_ST_ using a window size of 1,000. Pairwise *F*_ST_ values were then used to generate isolation by distance plots, using two different approaches for measuring distance between sampling sites: (i) seascape connectivity as calculated using the biophysics model, with connectivity data for the 64-generation model and averaged across all months, sampling depths, and drift durations, and (ii) shortest distance over the sea, calculated in R using the *shortestPath* function of the R-package gdistance v1.3.6 (88). We used a Manteltest to test for significant isolation by distance with the *mantel.rtest* function implemented in the R-package ade4 v1.7.19, using 9,999 repetitions (89).

We used PoPoolation v1.2.2 (70) to characterize genome-wide patterns of genetic variation and differentiation. To do so, we first obtained a pileup file for each pool using SAMtools mpileup with minimum base quality score and minimum alignment score of 20, and using the indel realigned BAM files as input. We used the resulting pileup files to calculate nucleotide diversity π and population mutation rate θ_W_ (Watterson’s theta) for each pool, using a minimum covered fraction of 20%, a window and step size of 1,000, minimum allele count of 4, minimum coverage of 20, and maximum coverage that equaled twice the sequencing depth across the genome. For Tajima’s D, we used the same settings, with the exceptions of a minimum allele count of 2 and a minimum coverage of 10. We used a t-test in R to test for significant differences in π, θ_W_, and Tajima’s D between the North Sea and Baltic Sea.

### Genotype-environment association (GEA) analyses for the Baltic Sea

Environmental data from the surface layer (0–10 m depth) of the North and Baltic seas for the period 2010–2018 were downloaded from ICES (https://www.ices.dk/data/data-portals/Pages/ocean.aspx) and Sharkweb (https://sharkweb.smhi.se/hamta-data/), and filtered to include environmental variables for which sufficient data were available to obtain a reliable interpolation across the study area. The final dataset included alkalinity, ammonium, dissolved oxygen, nitrate, nitrite, pH, phosphate, salinity, Secchi depth, silicate, temperature, total nitrogen, and total phosphorus. All data were interpolated across the study area using inverse distance weighting with the *idw* function of the R-package gstat v2.0.9 (90). For each sampling locality, we then extracted mean seasonal and annual values of each environmental variable from the interpolation. Although the main bloom period of *S. marinoi* in the Baltic Sea occurs between late February and early May, we chose to work with seasonal/annual values based on the consideration that *S. marinoi* also shows an autumn bloom and can be found year-round in the surface waters of the Baltic Sea (85).

We ran a PCA with the R-package vegan v2.6.4 (91) using the seasonal environmental variables and retaining informative axes using a broken stick model (PC1 and PC2). The PCA indicated that the North Sea (sample localities A and B) differed strongly from the Baltic Sea. Given that the SNP data of localities A and B show large *F*_ST_ differentiation with all samples in the Baltic Sea, this indicates that population structure and environmental variables are correlated. Univariate GEA methods do not perform well in such cases of distinct population differentiation (33, 92), so we ran GEA analyses on SNP data from the Baltic Sea localities only. We checked the environmental data of the Baltic for collinearity using the *removeCollinearity* function of the R-package virtualspecies v1.5.1 (93). If all seasonal measurements of the same variable had an R-squared value >0.7, the seasonal values were replaced by annual values. This was the case for salinity, pH, and ammonium. We ran a PCA on the resulting set of environmental variables, and retained informative axes using a broken stick model (PC1 and PC2). To detect associations between allele frequencies and environmental variables in the Baltic Sea, we ran two types of GEA analyses using the PC axis scores as representations of the environment, thus taking collinearity into account.

First, we ran LFMM using the *lfmm_ridge* model of the R-package lfmm v1.1 (35), using a K-value of 1. The latter was determined by running a PCA on the allele frequencies and assessing axis scores using a broken stick model. Analysis of Pool-seq data with LFMM can result in loss of power to detect outlier SNPs. To help alleviate this issue, we ran LFMM with a set of simulated ‘genotypes’ which for each pool equaled the number of strains/single cells that were originally included in the pool. Simulations were done using the R function *rbeta*, following the FAQ page of LFMM (http://membres-timc.imag.fr/Olivier.Francois/lfmm/faq.htm), and using a script adapted from the *rbeta_loop.R* script of (94). We performed GIF-correction of the P-values and subsequently controlled for multiple testing using an FDR <1%. To test whether the detected associations represent false positives, we randomized allele frequencies across the samples from the Baltic Sea. The randomized LFMM only detected 2 outlier SNPs for PC1 and none for PC2 with FDR <1%.

Second, we ran the auxiliary covariate (AUX) model in BayPass v2.3 (36). We first obtained a covariance matrix by running the core model in BayPass, using the default values (i.e., 20 pilot runs of 500 iterations, sampling every 25th iteration, and using a burnin of 5,000), with exception of the -d0yij parameter, which was set to 1/5th of the minimum pool size (i.e., 6.8). We ran the model three times, using three different seed values, and checked for convergence in R. We then used the covariance matrices obtained by the core model to run the AUX model. Using the same settings as the core model, we ran the AUX model three times using three different seed values, and each of the three covariance matrices from the three core model runs. The results of the runs were evaluated for convergence, after which we calculated the median Bayes factor across all three runs. We considered SNPs with a Bayes Factor (measured in deciban, dB) >20 as associated with the PC axes (36). Finally, we tested whether the overlap between outlier SNPs detected by LFMM and BayPass was larger than expected by chance using a hypergeometric test with the R-function *phyper*.

### Detecting outlier SNPs between the North Sea and the Baltic Sea

We identified outlier SNPs between the North Sea and the Baltic Sea using two approaches. To do so, we first combined all data of both seas by summarizing allele counts across both regions, effectively creating one observation for the North Sea and one for the Baltic Sea.

For the first approach, we calculated *F*_ST_ for each SNP in Poolfstat (69). In addition, we performed Fisher’s Exact Test using the *fisher-test.pl* script of PoPoolation2 on the full SNP dataset. P-values were corrected for multiple testing using Bonferroni correction. Outliers between the North Sea and the Baltic Sea were then defined as the top 1% outlier SNPs identified by *F*_ST_ that also showed a significant P-value in Fisher’s Exact Test after Bonferroni correction.

For the second approach, we ran the core model in BayPass v2.3 (36), using the default values (20 pilot runs of 500 iterations, sampling every 25th iteration, and using a burnin of 5,000), with the exception of the -d0yij parameter, which was set to 1/5th of the minimum pool size (i.e., 16.2, thus taking into account that allele counts were summarized across the North Sea and the Baltic Sea), following instructions in the manual. We ran the model three times, using three different seed values, and checked for convergence in R. BayPass outliers were then detected using two approaches. First, we generated three pseudo-observation datasets (PODs) of 100,000 SNPs and reran BayPass using the PODs and the same settings as used for the core model. The PODs output was then evaluated against the original core models, and outliers were defined as all SNPs located above the 99% significance threshold of the subpopulation differentiation *XtX* identified using the PODs, following instructions in the manual. Outlier SNPs were only accepted if identified by each of the three model runs. Second, we used FDR-control of the P-values generated by BayPass to detect outliers, using an FDR-threshold of 1%. Finally, we combined the outliers of the POD and FDR approaches, and only retained those that were identified by both approaches.

### Functional annotation and GO enrichment of outlier genes

We obtained functional annotations of the complete set of genes in the *S. marinoi* reference genome v1.1.2. First, we used BLAST+ v2.6.0 (95) to run sequence similarity blastp searches of all *S. marinoi* proteins annotated in the genome against the Swissprot (download September 2022) and Uniprot databases (download June 2019), and retained the best hit using a maximum e-value limit of 1e-6. Second, we ran InterProScan v5.36-75.0 (96) against the InterPro collection of protein signature databases, including GO resources, Pfam domains, PRINTS, PANTHER, SMART, SignalP_EUK, and TMHMM. Third, we obtained KEGG pathway annotations via the KofamKOALA web server v2022-08-01 (KEGG release 103.0) (97).

The sync file with all SNPs that passed quality control was converted into a VCF file with the script *sync2VCF.R*, and SNPs were subsequently annotated with SnpEff v5.1 (71). For each outlier SNP, we used SnpEff annotations to identify whether the SNP was located in an exon, UTR region, intron, or intergenic region. When a SNP was located in an exon or UTR region, we identified its corresponding gene. In several cases, a SNP was positioned in two genes located on opposite strands: in these cases both genes were considered to be affected by the SNP. In some cases, the allele of the reference genome was not detected as a SNP in our dataset, resulting in two alternative alleles. In these cases, we assumed that transversions correspond with missense SNPs, whereas transitions correspond with synonymous SNPs. Using the resulting list of genes with outlier SNPs (outlier genes), we performed GO enrichment on the *F*_ST_-outlier and GEA-outlier genes, distinguishing between the full set of outlier genes, and outlier genes with missense SNPs, SNPs in UTR regions, or outlier genes with multiple SNPs. This was done using Fisher’s Exact Test and the *elim* algorithm in the R-package TopGO v2.46.0 (72), separately for each main GO category (biological process, molecular function, and cellular component). The background gene set of GO terms represented the full set of GO terms extracted from the *S. marinoi* reference genome v1.1.2.

### Linking outlier SNPs to gene expression patterns

To link the outlier genes with previously published salinity transcriptomes of *S. marinoi*, we first reanalyzed the transcriptome data using the *S. marinoi* reference genome v1.1.2, given that in the original study we used an older version of the genome (v1.1) (11). This reanalysis detected virtually the same set of differentially expressed genes, but found a larger number of ‘core response genes’ deemed essential for low salinity acclimation in *S. marinoi* (33 vs. 27) (11). Using the updated transcriptome data, we tested whether the overlap in differentially expressed genes and *F*_ST_-outlier or GEA-outlier genes was significant using the R function *phyper*. The reference set for the significance test included all 17,203 genes in the genome of *S. marinoi*. The differentially expressed genes were separated into two major sets: (i) genes that were differentially expressed in response to salinity in at least one of the eight investigated strains and/or the average response of all eight strains examined together (average and strain-specific response), and (ii) genes that showed significantly different gene expression responses in different strains (interaction-effects). We visually explored the relationship between the magnitude of gene expression differences between strains (interaction-effects) and between-pool allele frequency differences of SNPs within corresponding genes.

## DATA AVAILABILITY

Pool-seq data are available from the Sequence Read Archive (NCBI) under project number PRJNA950465. The *S. marinoi* reference genome (v1.1.2) used for read mapping and reanalysis of the *S. marinoi* salinity transcriptome, as well as all datasets and scripts needed to reproduce the analyses and figures are available from Zenodo (10.5281/zenodo.7786015).

## AUTHOR CONTRIBUTIONS

AJA and AG conceived the project. AJA, ECR, TN, AG, and MWH designed the study. OK, AK, CS, MT, and AG performed fieldwork and aided with experimental design. AG, MT, and MIMP provided the annotated genome of *S. marinoi*. ECR performed laboratory work, including culturing and molecular work, with support from TN. TN wrote the machine learning script for estimating the number of cells per chain. PRJ designed and ran the biophysics model for seascape connectivity. EP conceived the data analysis workflow and analyzed the data, with support from MWH, WRR, and TN. EP wrote the manuscript and prepared the figures. All authors read and commented on the manuscript.

## ACKNOWLEDGEMENTS

This work was supported by grants from the Simons Foundation (403249, AJA and 725407, EP) and a grant from the National Science Foundation (1651087, AJA). This research used resources available through the Arkansas High Performance Computing Center, which is funded through multiple NSF grants and the Arkansas Economic Development Commission. We thank Geoffrey House for help with the sampling design, and we are grateful to Sirje Sildever (Tallinn University of Technology, Estonia), Björn Andersson (University of Gothenburg, Sweden), Andrzej Witkowski (University of Szczecin, Poland), Jörg Dutz (Leibniz Institute for Baltic Sea Research Warnemuende, Germany), Justyna Kobos (University of Gdansk, Poland), and Anu Vehmaa (University of Helsinki, Finland) for sample collection.

## SUPPLEMENTARY TABLES

**Table S1.** | Sampling localities for population genomics. **a.** Sampling localities from which *S. marinoi* strains were obtained, showing the total number (#) of strains and cells included for each locality. *M* refers to ‘million’. Note that the sequences of strains from samples P1 and P2 were merged together after initial data processing.

| Pool | Collection | # Strains | # Cells | Material | Country | Region | GPS (N/E) | Collector | Year |
| --- | --- | --- | --- | --- | --- | --- | --- | --- | --- |
| A | AJA304 | single cell | 41 | Sediment | Sweden | North Sea (near Gothenburg) | 58.25000/11.05000 | A.Godhe | 2014 |
| B | AJA305 | 40 | 40M | Sediment | Sweden | Danish Straits (Øresund), near Malmö | 55.97744/12.69058 | A.Godhe | 2010 |
| D | AJA332 | single cell | 40 | Sediment | Sweden | Baltic Sea, Gropviken | 58.33200/16.70583 | A.Godhe, B.Andersson | 2017 |
| F | AJA328 | 41 | 40M | Sediment | Sweden | Baltic Sea, Gulf of Bothnia | 62.65317/18.95200 | A.Godhe | 2016/2017 |
| H | AJA331 | single cell | 40 | Sediment | Finland | Baltic Sea, Archipelago Sea, Tvärminne | 59.88222/23.25389 | A.Kremp | 2018 |
| I | AJA311 | 40 | 40M | Sediment | Finland | Baltic Sea, Gulf of Finland | 60.18000/25.50700 | A.Kremp | 2015 |
| J | AJA313 | 39 | 25M | Sediment | Finland | Baltic Sea, Gulf of Finland, near Loviisa | 60.37833/26.36525 | A.Kremp | 2016 |
| K | AJA318 | 34 | 40M | Sediment | Estonia | Baltic Sea, Southeast coast near Irbe strait | 57.81670/22.28330 | S.Sildever | 2016 |
| P1 | AJA334 | 33 | 40M | Plankton | Poland | Baltic Sea, south coast | 54.44778/18.57611 | A.Witkowski | 2018 |
| P2 | AJA333 | 18 | 40M | Sediment | Poland | Baltic Sea, south coast | 54.44778/18.57611 | A.Witkowski | 2018 |

## SUPPLEMENTARY FIGURES

**Supplementary Fig. S1.**
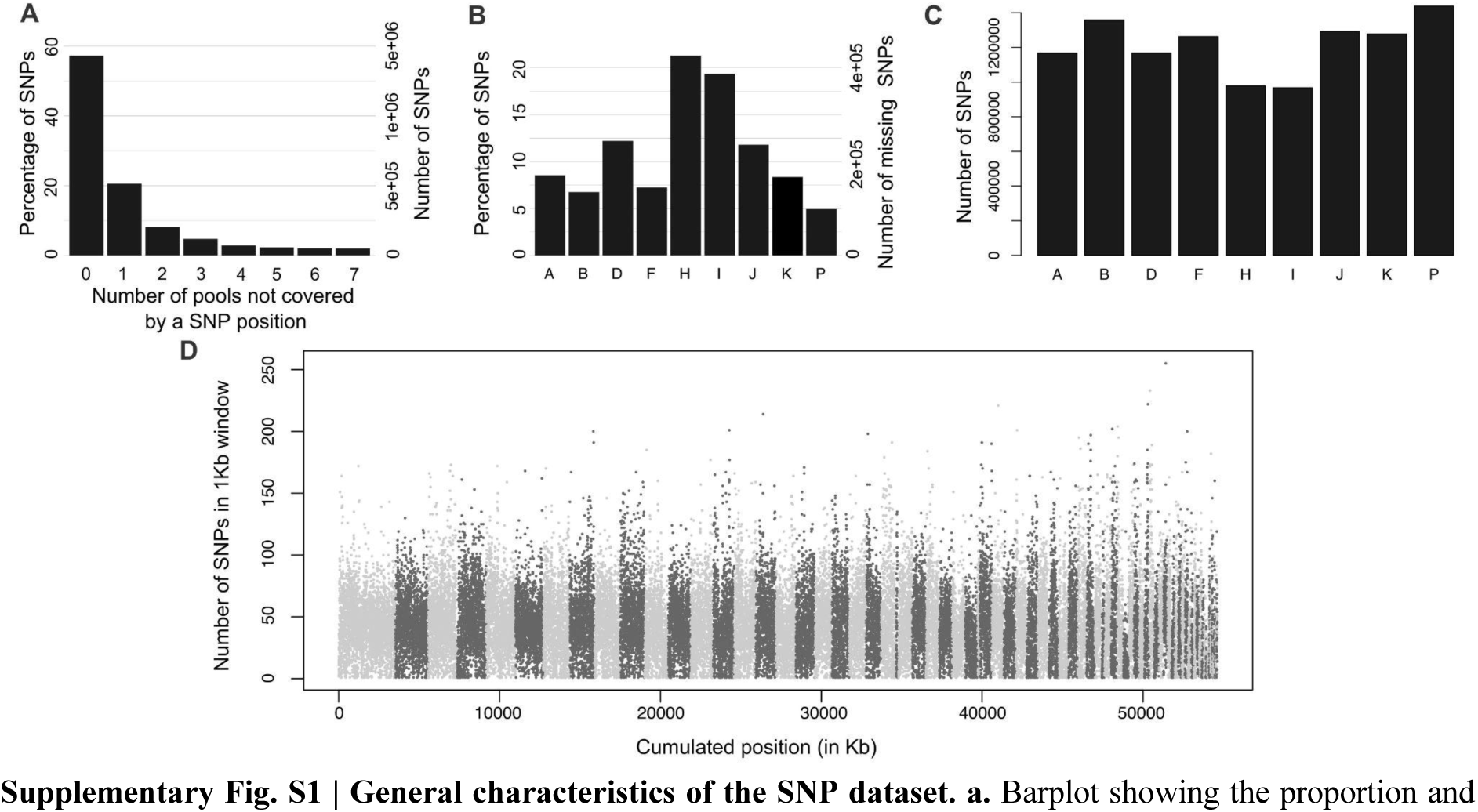
| General characteristics of the SNP dataset. **a.** Barplot showing the proportion and number of SNPs that are missing *n* times from a pool. The bar at zero indicates the total proportion/number of SNPs that were covered in each position. **b.** Barplot showing the proportion and number of ‘missing’ SNPs for each pool. A ‘missing SNP’ is defined as a SNP that was not covered in at least one pool, for example due to insufficient coverage. **c.** Number of SNPs within each pool, calculated from the set of 1,259,039 SNP positions that were covered in each pool. **d.** Manhattan plot showing the distribution of the SNPs across the reference genome of *S. marinoi*. Different contigs are indicated by the alternating shades of gray.

**Supplementary Fig. S2.**
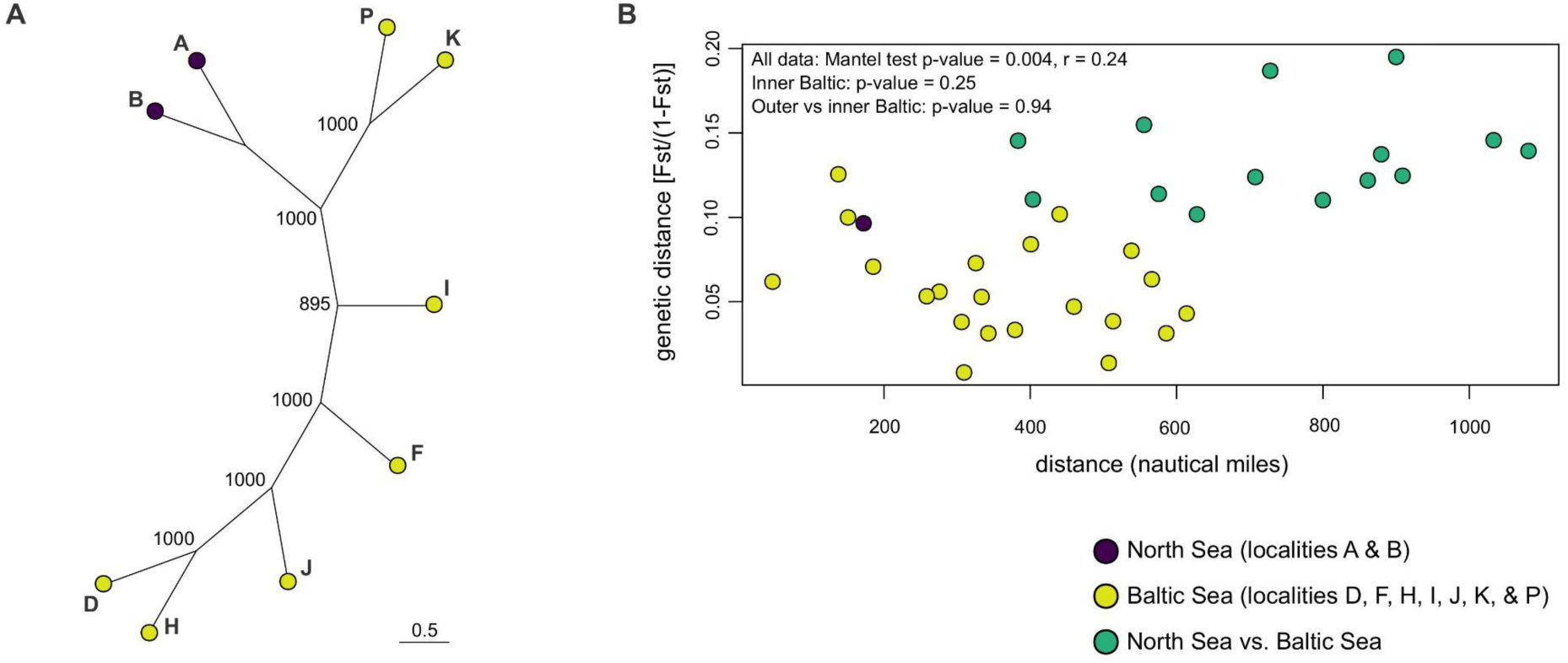
| Population structure of *S. marinoi*. **a.** Neighbor-joining tree of the allele frequencies, showing bootstrap support values. **b.** Isolation-by-distance plot. Distance is measured as the shortest distance over the sea, in nautical miles. Prior to creating the plots in panels a-b, all SNPs identified as outliers were removed.

**Supplementary Fig. S3.**
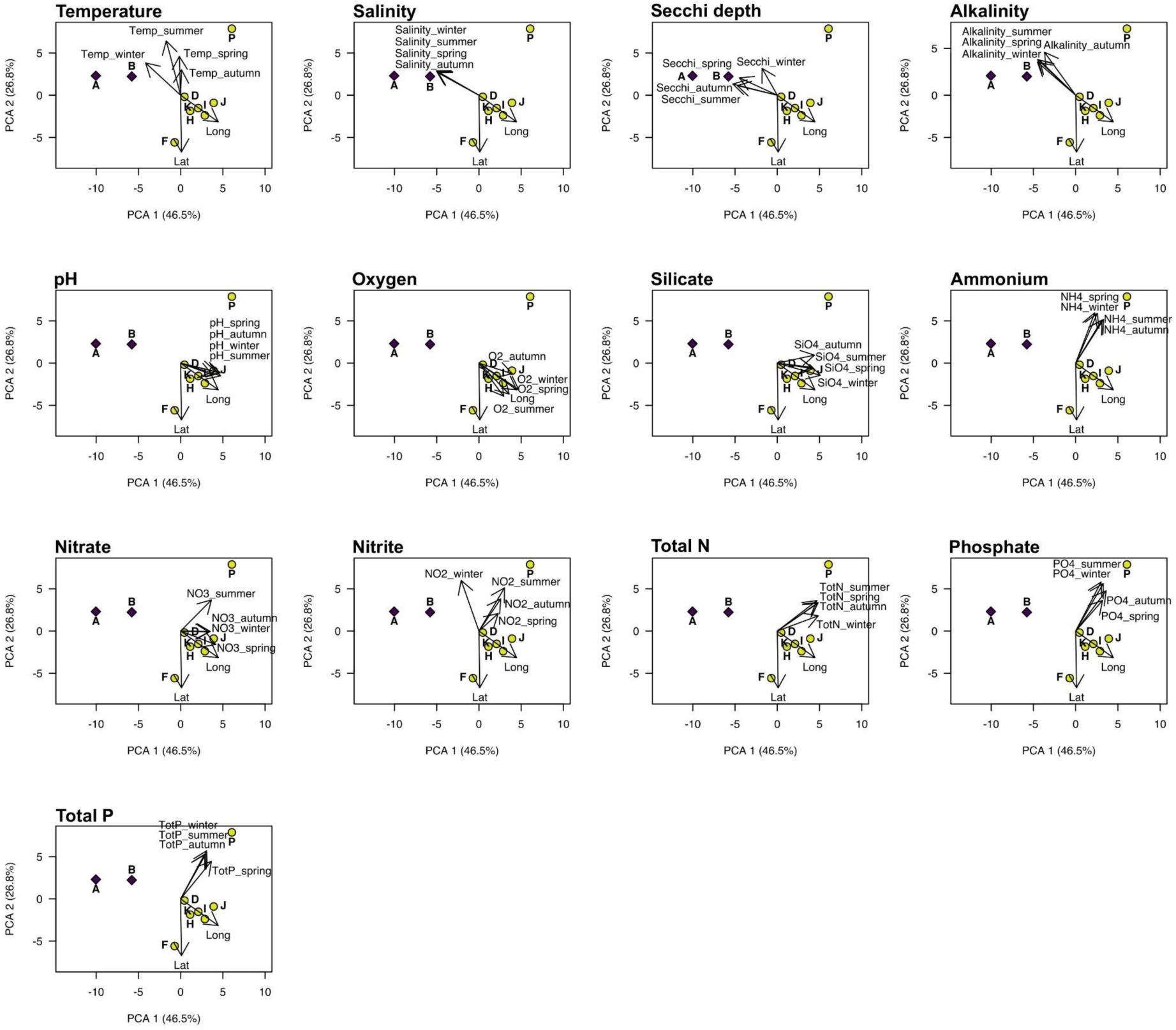
| The abiotic environment of the North Sea and the Baltic Sea. Principal Component Analysis (PCA) plots on the environmental variables from the North Sea and Baltic Sea. Each plot represents the same analysis/PC axes, but with indication of different environmental variables, for sake of clarity. The following environmental variables were included: alkalinity, ammonium (NH_4_), dissolved oxygen (O_2_), nitrate (NO_3_), nitrite (NO_2_), pH, phosphate (PO_4_), salinity, secchi depth, silicate (SiO_4_), temperature (temp), total nitrogen (totN), and total phosphorus (totP). Only PC axes 1 and 2 are shown, as the broken stick model indicated the other axes were not informative.

**Supplementary Fig. S4.**
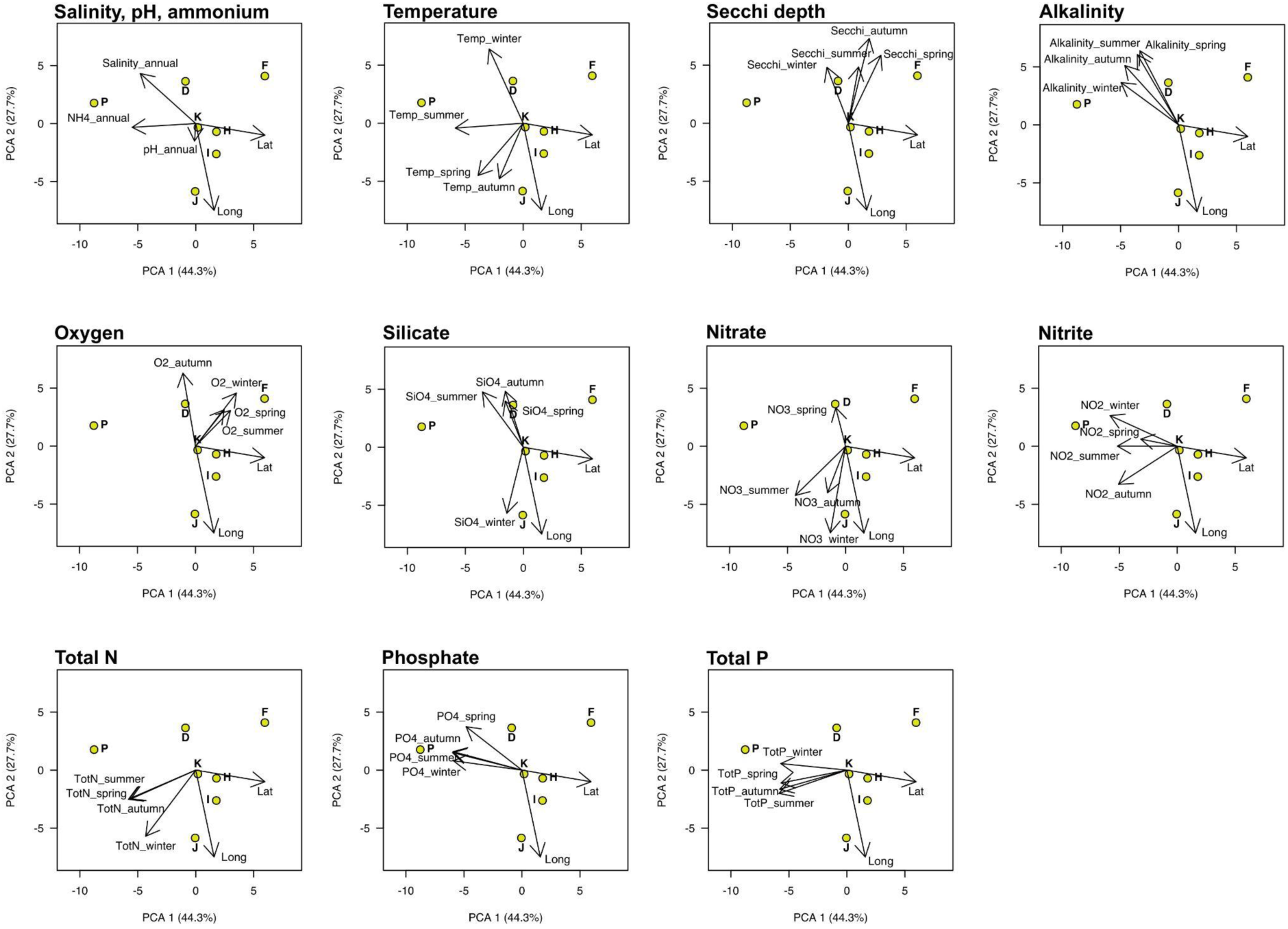
| The abiotic environment of the Baltic Sea. Principal Component Analysis (PCA) plots on the environmental variables from the Baltic Sea. Each plot represents the same analysis/PC axes, but with indication of different environmental variables, for sake of clarity. The following environmental variables were included: alkalinity, ammonium (NH_4_), dissolved oxygen (O_2_), nitrate (NO_3_), nitrite (NO_2_), pH, phosphate (PO_4_), salinity, secchi depth, silicate (SiO_4_), temperature (temp), total nitrogen (totN), and total phosphorus (totP). Only PC axes 1 and 2 are shown, as the broken stick model indicated the other axes were not informative.

**Supplementary Fig. S5.**
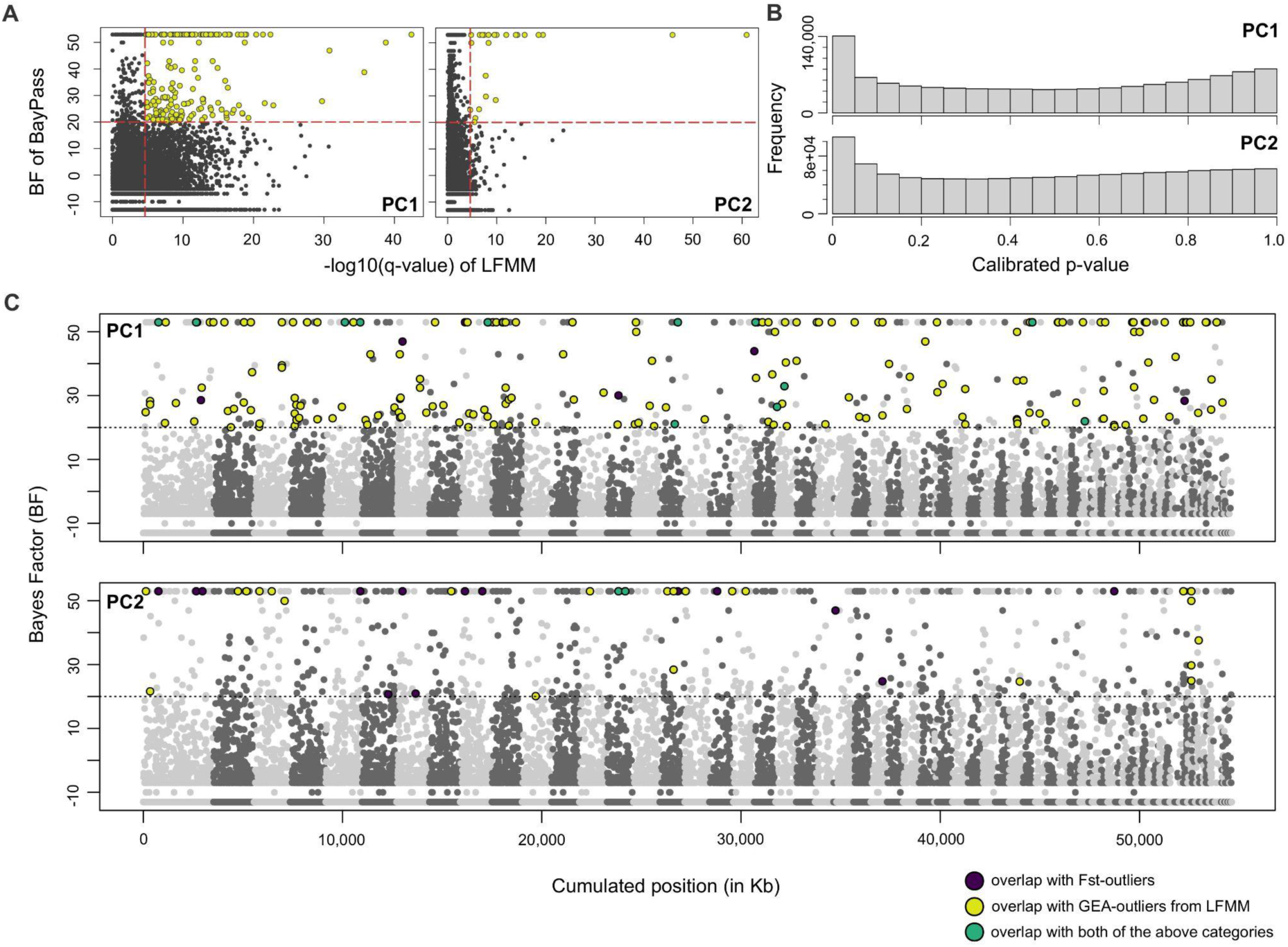
| Outlier SNPs associated with the Baltic Sea environmental gradients. **a.** Dotplots showing overlap of outlier SNPs detected by LFMM and BayPass AUX for PC1 (north-south gradient) and PC2 (east-west gradient). The vertical and horizontal red lines indicate the significance thresholds for LFMM (FDR < 1%) and BayPass (BF > 20), respectively. SNPs indicated in yellow were found to be significant outliers for both LFMM and BayPass AUX. **b.** LFMM p-value distributions for PC1 and PC2 after GIF correction (GIF PC1 = 25.8, GIF PC2 = 31.3). **c.** Manhattan plots showing the outlier SNPs that are associated with the environment of the Baltic Sea as estimated by BayPass, shown separately for both PC axes. The dotted line represents the 20 BF threshold above which SNPs are considered to be significantly associated with the environment. Different contigs are indicated with alternating shades of gray. The colored dots represent outlier SNPs that overlap with the *F*_ST_-outliers (dark blue), LFMMs GEA test (yellow), and both of the former (green).

**Supplementary Fig. S6.**
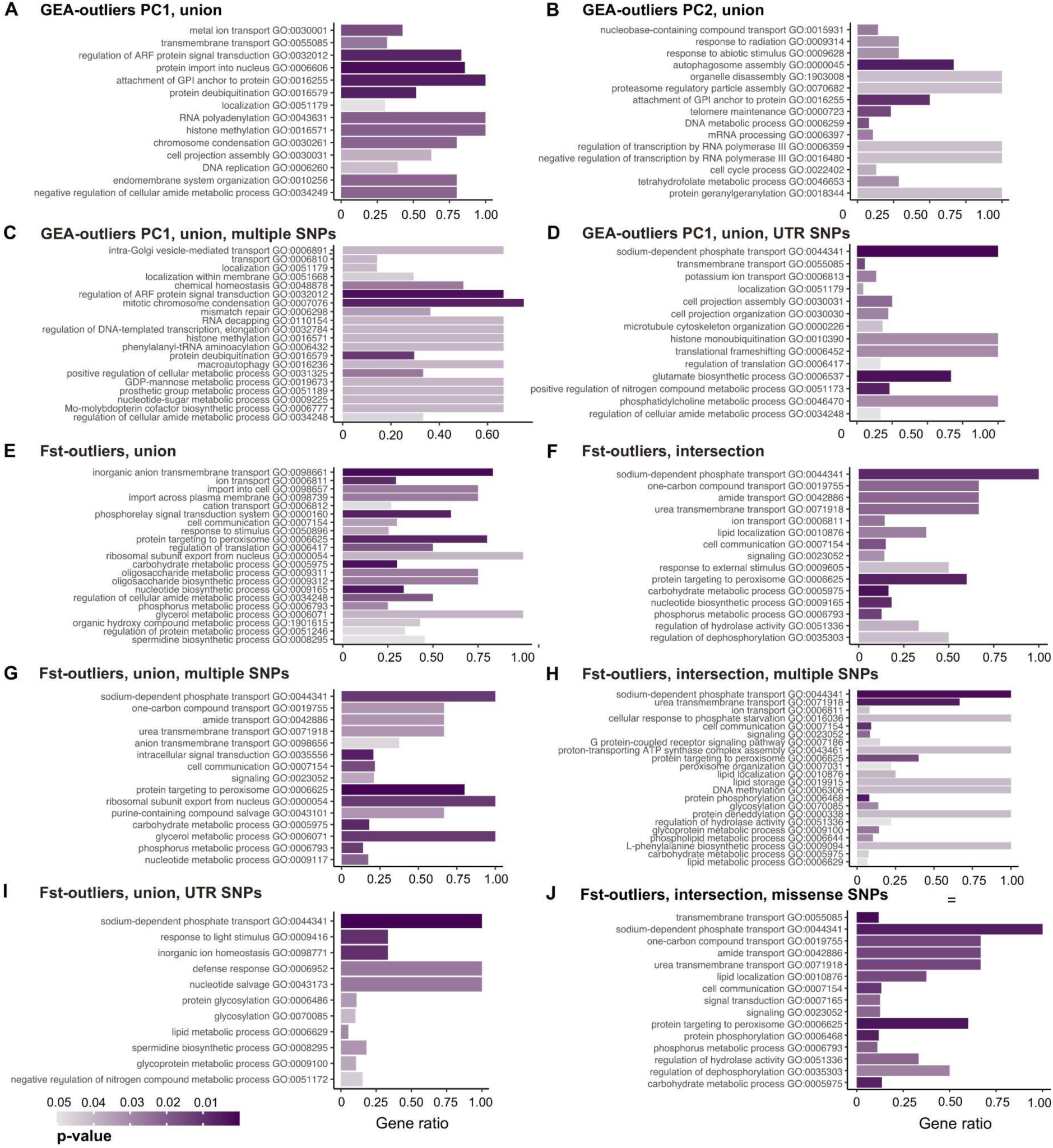
| Gene Ontology (GO) enrichment of outlier genes. The barplots show GO enrichment results for several sets of outlier genes. **a-d.** GEA-outlier genes: union of both approaches (LFMM/BayPass) for PC1 (north-south gradient) (**a**) and PC2 (east-west gradient) (**b**), PC1 GEA-outlier genes only including genes that have more than one outlier SNP (union of both approaches) (**c**), PC1 GEA-outlier genes only including genes that have SNPs in the UTR regions (union of both approaches) (**d**). **e-j.** *F*_ST_-outlier genes: union (**e**) and intersection (**f**) on all outlier SNPs identified by both approaches (BayPass/FET), *F*_ST_-outliers only including genes with multiple SNPs, showing the union (**g**) and intersection (**h**) of both approaches, *F*_ST_-outlier genes only including genes with SNPs in the UTR regions (union of both approaches) (**i**), and *F*_ST_-outlier genes only including genes with missense SNPs (intersection of both approaches) (**j**). Bars are colored by topGO’s p-value. For each plot, GO terms are sorted by general theme, and only include Biological Process GO terms that were retained after running REVIGO. The height of the bars indicates the proportion of genes with a given GO term that are enriched relative to the total number of genes with this GO term in the genome of *S. marinoi*.

**Supplementary Fig. S7.**
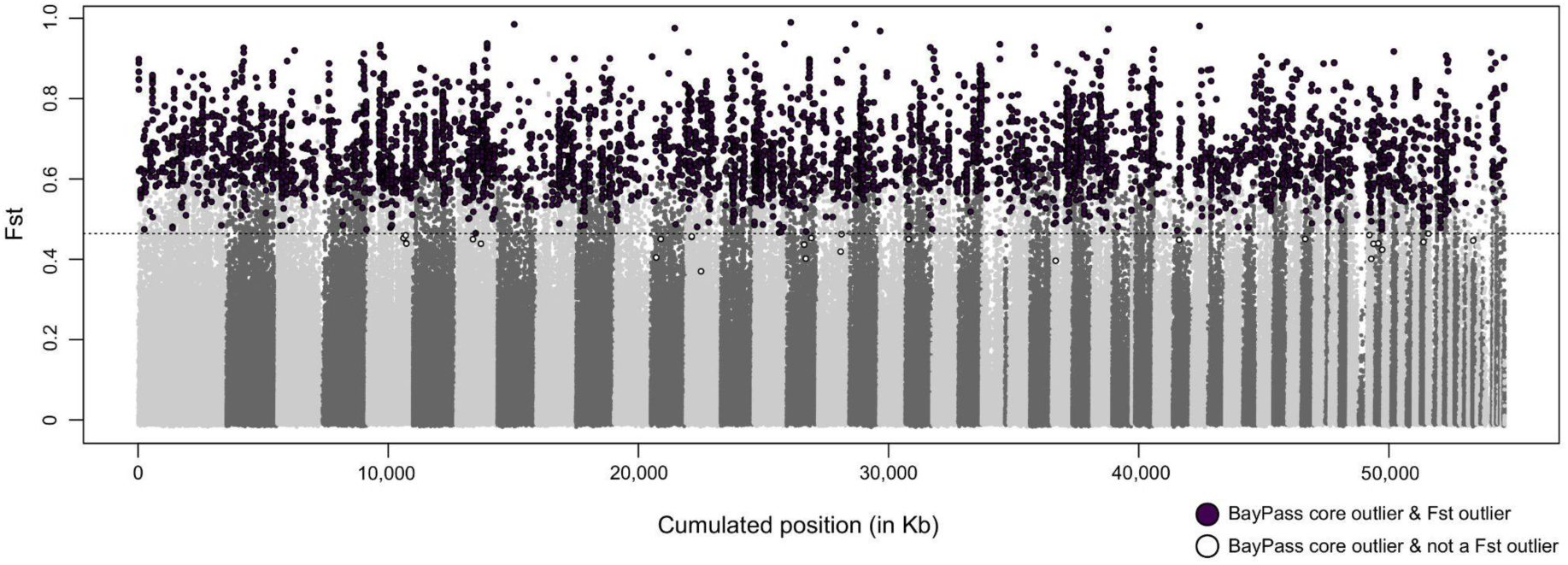
| *F*_ST_-Outlier SNPs contrasting the North Sea and the Baltic Sea. Manhattan plot showing the *F*_ST_-outlier SNPs, generated by contrasting the North Sea and the Baltic Sea localities. The plot shows *F*_ST_-outliers, which were defined as the top 1% SNPs with highest *F*_ST_ scores that also showed a significant P-value in Fisher’s Exact Test (FET) (note that all SNPs above the 1% threshold were significant). The dotted line represents the top 1% *F*_ST_ threshold. The purple dots represent *F*_ST_-outlier SNPs that overlap with the outlier SNPs detected by the BayPass core model. The white dots represent the outlier SNPs detected by the BayPass core model that were not detected by FET.

**Supplementary Fig. S8.**
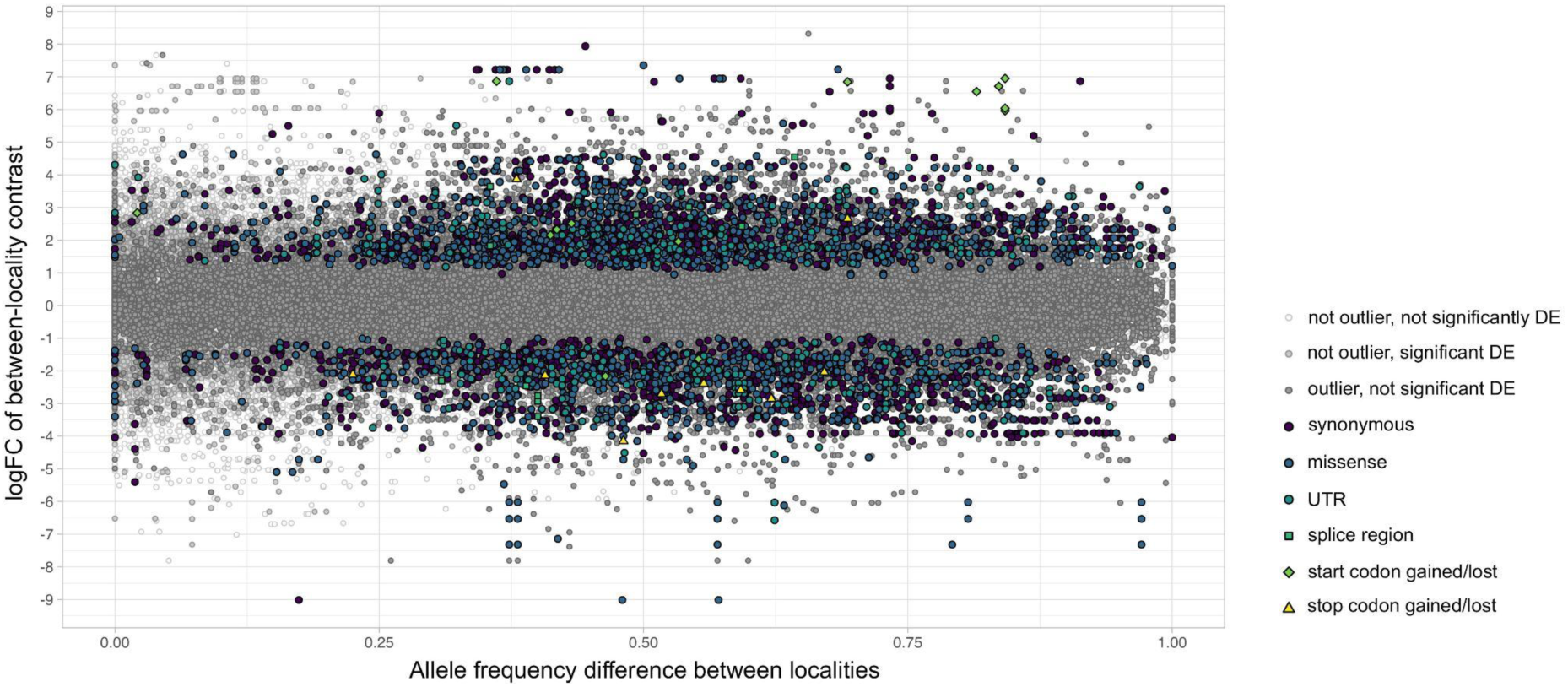
| Gene expression differences in function of allele frequency differences. The scatterplot shows the link between allele frequency differences between localities and gene expression differences (logFC) in response to low salinity between strains from the same localities. Gene expression data were reanalyzed from (11), using the same version of the *S. marinoi* genome used for read mapping in this study. Gene expression data were obtained from eight strains, originating from localities A, B, D, F, I, J, K, and P, which were exposed to three different salinities mimicking the Baltic Sea salinity cline (24, 16, and 8). For this reason, data from locality H are not included in the plot. For all combinations of the eight strains, we tested for interaction-effects, thus revealing significant differences in the response between strains to low salinity. This resulted in a logFC value for each gene for each strain pair. Only logFC values of salinity 8 vs. salinity 24 were considered for the plot. Allele frequency differences were calculated by taking the absolute value of the frequency difference of the major allele for each pair of localities. We then linked each SNP with its corresponding gene, and the logFC of between-locality contrasts. Since there are multiple logFC values for each gene (and thus allele frequency difference), allele frequencies are repeated in the plot across pairs of localities. If a gene had multiple SNPs, the same logFC value was linked to all relevant allele frequency differences. In the plot, gray dots indicate SNPs that were not outliers and/or that were not differentially expressed (DE) in response to salinity. Colored dots are SNPs that were differentially expressed between strains in response to salinity, and which were also identified as outlier SNPs. The colors and shape of the symbols indicate the type of outlier SNPs: synonymous mutations, missense mutations, SNPs located in the UTR regions or splice regions, and SNPs that resulted in the gain/loss of a start or stop codon.

**Supplementary Fig. S9.**
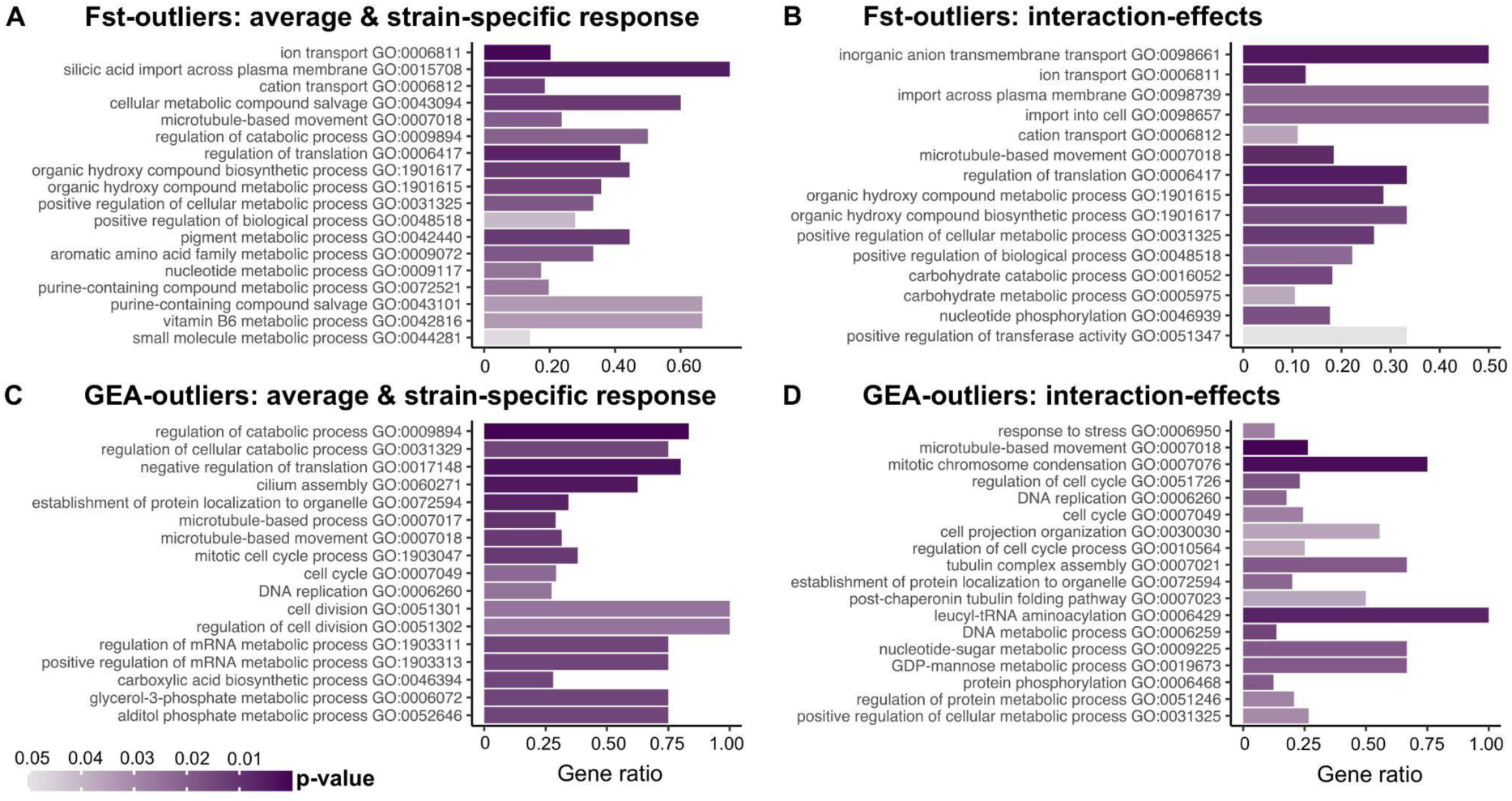
| GO enrichment on the overlap between outlier and differentially expressed genes. Barplots showing GO enrichment results for the overlap between outlier genes and differentially expressed genes: overlap between *F*_ST_-outlier genes (union of both approaches) and the average & strain-specific response of the RNA-seq (**a**) and the interaction-effects (**b**), overlap between GEA-outlier genes (union of both approaches) and the average & strain-specific response of the RNA-seq (**c**) and the interaction-effects (**d**). Bars are colored by topGO’s P-value. For each plot, GO terms are sorted by general theme, and only include Biological Process GO terms that were retained after running REVIGO. The height of the bars indicates the proportion of genes with a given GO term that are enriched relative to the total number of genes with this GO term in the genome of *S. marinoi*.

## REFERENCES

1. A. Z. Worden, et al., Environmental science. Rethinking the marine carbon cycle: factoring in the multifarious lifestyles of microbes. Science 347, 1257594 (2015).

2. R. Cavicchioli, et al., Scientists’ warning to humanity: microorganisms and climate change. Nat. Rev. Microbiol. 17, 569–586 (2019).

3. L. Schlüter, K. T. Lohbeck, J. P. Gröger, U. Riebesell, T. B. H. Reusch, Long-term dynamics of adaptive evolution in a globally important phytoplankton species to ocean acidification. Sci Adv 2, e1501660 (2016).

4. D. Padfield, G. Yvon-Durocher, A. Buckling, S. Jennings, G. Yvon-Durocher, Rapid evolution of metabolic traits explains thermal adaptation in phytoplankton. Ecol. Lett. 19, 133–142 (2016).

5. D. A. Filatov, How does speciation in marine plankton work? Trends Microbiol., 10.1016/j.tim.2023.07.005 (2023).

6. F. Moerman, E. A. Fronhofer, F. Altermatt, A. Wagner, Selection on growth rate and local adaptation drive genomic adaptation during experimental range expansions in the protist *Tetrahymena thermophila*. J. Anim. Ecol. 91, 1088–1103 (2022).

7. C.-E. Schaum, A. Buckling, N. Smirnoff, D. J. Studholme, G. Yvon-Durocher, Environmental fluctuations accelerate molecular evolution of thermal tolerance in a marine diatom. Nat. Commun. 9, 1719 (2018).

8. K. E. Helliwell, et al., Fundamental shift in vitamin B12 eco-physiology of a model alga demonstrated by experimental evolution. ISME J. 9, 1446–1455 (2015).

9. A. Rastogi, et al., A genomics approach reveals the global genetic polymorphism, structure, and functional diversity of ten accessions of the marine model diatom *Phaeodactylum tricornutum*. ISME J. 14, 347–363 (2020).

10. P. Bulankova, et al., Mitotic recombination between homologous chromosomes drives genomic diversity in diatoms. Curr. Biol. 31, 3221–3232.e9 (2021).

11. E. Pinseel, et al., Strain-specific transcriptional responses overshadow salinity effects in a marine diatom sampled along the Baltic Sea salinity cline. ISME J. 16, 1776–1787 (2022).

12. E. Schaum, B. Rost, A. J. Millar, S. Collins, Variation in plastic responses of a globally distributed picoplankton species to ocean acidification. Nat. Clim. Chang. 3, 298–302 (2012).

13. I. W. Bishop, S. I. Anderson, S. Collins, T. A. Rynearson, Thermal trait variation may buffer Southern Ocean phytoplankton from anthropogenic warming. Glob. Chang. Biol. 28, 5755–5767 (2022).

14. M. Aranguren-Gassis, C. T. Kremer, C. A. Klausmeier, E. Litchman, Nitrogen limitation inhibits marine diatom adaptation to high temperatures. Ecol. Lett. 22, 1860–1869 (2019).

15. J. Zhong, et al., Adaptation of a marine diatom to ocean acidification and warming reveals constraints and trade-offs. Sci. Total Environ. 771, 145167 (2021).

16. C. Nef, M.-A. Madoui, É. Pelletier, C. Bowler, Whole-genome scanning reveals environmental selection mechanisms that shape diversity in populations of the epipelagic diatom *Chaetoceros*. PLoS Biol. 20, e3001893 (2022).

17. S. Björck, A review of the history of the Baltic Sea, 13.0-8.0 ka BP. Quat. Int. 27, 19–40 (1995).

18. C. A. Lozupone, R. Knight, Global patterns in bacterial diversity. Proceedings of the National Academy of Sciences 104, 11436–11440 (2007).

19. F. van Wirdum, et al., Middle to Late Holocene variations in salinity and primary productivity in the central Baltic Sea: a multiproxy study from the Landsort Deep. Frontiers in Marine Science 6, 51 (2019).

20. A. Godhe, et al., Physical barriers and environmental gradients cause spatial and temporal genetic differentiation of an extensive algal bloom. J. Biogeogr. 43, 1130–1142 (2016).

21. C. Sjöqvist, A. Godhe, P. R. Jonsson, L. Sundqvist, A. Kremp, Local adaptation and oceanographic connectivity patterns explain genetic differentiation of a marine diatom across the North Sea-Baltic Sea salinity gradient. Mol. Ecol. 24, 2871–2885 (2015).

22. K. Johannesson, A. Le Moan, S. Perini, C. André, A Darwinian laboratory of multiple contact zones. Trends Ecol. Evol. 35, 1021–1036 (2020).

23. K. Rengefors, et al., Genome-wide single nucleotide polymorphism markers reveal population structure and dispersal direction of an expanding nuisance algal bloom species. Mol. Ecol. 30, 912–925 (2021).

24. A. Godhe, K. Härnström, Linking the planktonic and benthic habitat: genetic structure of the marine diatom Skeletonema marinoi. Mol. Ecol. 19, 4478–4490 (2010).

25. K. Härnström, M. Ellegaard, T. J. Andersen, A. Godhe, Hundred years of genetic structure in a sediment revived diatom population. Proceedings of the National Academy of Sciences 108, 4252–4257 (2011).

26. D. B. Lowry, et al., Breaking RAD: an evaluation of the utility of restriction site-associated DNA sequencing for genome scans of adaptation. Mol. Ecol. Resour. 17, 142–152 (2017).

27. R. Gollnisch, J. Wallenius, K. E. Gribble, D. Ahrén, K. Rengefors, SAG-RAD: a method for single-cell population genomics of unicellular eukaryotes. Mol. Biol. Evol. msad095 (2023).

28. M. Slimp, L. D. Williams, H. Hale, M. G. Johnson, On the potential of Angiosperms353 for population genomic studies. Appl. Plant Sci. 9 (2021).

29. C. White, et al., Ocean currents help explain population genetic structure. Proc. Biol. Sci. 277, 1685–1694 (2010).

30. M. Jahnke, et al., Seascape genetics and biophysical connectivity modelling support conservation of the seagrass *Zostera marina* in the Skagerrak-Kattegat region of the eastern North Sea. Evol. Appl. 11, 645–661 (2018).

31. A. Godhe, T. Rynearson, The role of intraspecific variation in the ecological and evolutionary success of diatoms in changing environments. Philos. Trans. R. Soc. Lond. B Biol. Sci. 372, 20160399 (2017).

32. E. M. Leffler, et al., Revisiting an old riddle: what determines genetic diversity levels within species? PLoS Biol. 10, e1001388 (2012).

33. B. R. Forester, M. R. Jones, S. Joost, E. L. Landguth, J. R. Lasky, Detecting spatial genetic signatures of local adaptation in heterogeneous landscapes. Mol. Ecol. 25, 104–120 (2016).

34. K. E. Lotterhos, The paradox of adaptive trait clines with nonclinal patterns in the underlying genes. Proc. Natl. Acad. Sci. U. S. A. 120, e2220313120 (2023).

35. B. Jumentier, lfmm: Latent Factor Mixed Models. R package version 1.1. https://CRAN.R-project.org/package=lfmm (2021).

36. M. Gautier, Genome-wide scan for adaptive divergence and association with population-specific covariates. Genetics 201, 1555–1579 (2015).

37. M. Granskog, H. Kaartokallio, H. Kuosa, D. N. Thomas, J. Vainio, Sea ice in the Baltic Sea – A review. Estuar. Coast. Shelf Sci. 70, 145–160 (2006).

38. F. Mignone, C. Gissi, S. Liuni, G. Pesole, Untranslated regions of mRNAs. Genome Biol. 3, reviews0004.1 (2002).

39. A. Platt, P. F. Gugger, M. Pellegrini, V. L. Sork, Genome-wide signature of local adaptation linked to variable CpG methylation in oak populations. Mol. Ecol. 24, 3823–3830 (2015).

40. M. J. Dubin, et al., DNA methylation in *Arabidopsis* has a genetic basis and shows evidence of local adaptation. Elife 4, e05255 (2015).

41. I. Kaczmarska, et al., Proposals for a terminology for diatom sexual reproduction, auxospores and resting stages. Diatom Res. 28, 263–294 (2013).

42. A. Godhe, A. Kremp, M. Montresor, Genetic and microscopic evidence for sexual reproduction in the centric diatom *Skeletonema marinoi*. Protist 165, 401–416 (2014).

43. J. Sefbom, et al., Local adaptation through countergradient selection in northern populations of *Skeletonema marinoi*. Evol. Appl. 16, 311–320 (2023).

44. T. Berglund, A. B. Ohlsson, Defensive and secondary metabolism in plant tissue cultures, with special reference to nicotinamide, glutathione and oxidative stress. Plant Cell Tissue Organ Cult. 43, 137–145 (1995).

45. K. M. Downey, K. J. Judy, E. Pinseel, A. J. Alverson, J. A. Lewis, The dynamic response to hypo-osmotic stress reveals distinct stages of freshwater acclimation by a euryhaline diatom. Mol. Ecol. 32, 2766–2783 (2023).

46. A. Amato, et al., Grazer-induced transcriptomic and metabolomic response of the chain-forming diatom *Skeletonema marinoi*. ISME J. 12, 1594–1604 (2018).

47. J. Bergkvist, P. Thor, H. H. Jakobsen, S.-Å. Wängberg, E. Selander, Grazer-induced chain length plasticity reduces grazing risk in a marine diatom. Limnol. Oceanogr. 57, 318–324 (2012).

48. I. Vuorinen, J. Hänninen, M. Viitasalo, U. Helminen, H. Kuosa, Proportion of copepod biomass declines with decreasing salinity in the Baltic Sea. ICES J. Mar. Sci. 55, 767–774 (1998).

49. E. Selander, et al., Copepods drive large-scale trait-mediated effects in marine plankton. Sci Adv 5, eaat5096 (2019).

50. J. Sefbom, et al., A planktonic diatom displays genetic structure over small spatial scales. Environ. Microbiol. 20, 2783–2795 (2018).

51. J. Sefbom, I. Sassenhagen, K. Rengefors, A. Godhe, Priority effects in a planktonic bloom-forming marine diatom. Biol. Lett. 11, 20150184 (2015).

52. S. Sildever, J. Sefbom, I. Lips, A. Godhe, Competitive advantage and higher fitness in native populations of genetically structured planktonic diatoms. Environ. Microbiol. 18, 4403–4411 (2016).

53. L. Sundqvist, A. Godhe, P. R. Jonsson, J. Sefbom, The anchoring effect-long-term dormancy and genetic population structure. ISME J. 12, 2929–2941 (2018).

54. K. E. Helliwell, et al., Spatiotemporal patterns of intracellular Ca2+ signalling govern hypo-osmotic stress resilience in marine diatoms. New Phytol. 230, 155–170 (2021).

55. Z. Chen, et al., Phosphoproteomic analysis provides novel insights into stress responses in *Phaeodactylum tricornutum*, a model diatom. J. Proteome Res. 13, 2511–2523 (2014).

56. A. Pelusi, et al., Gene expression during the formation of resting spores induced by nitrogen starvation in the marine diatom *Chaetoceros socialis*. BMC Genomics 24, 106 (2023).

57. Q. Shen, et al., Dual activities of plant cGMP-dependent protein kinase and its roles in gibberellin signaling and salt stress. Plant Cell 31, 3073–3091 (2019).

58. S. Suzuki, et al., Rapid transcriptomic and physiological changes in the freshwater pennate diatom *Mayamaea pseudoterrestris* in response to copper exposure. DNA Res. 29 (2022).

59. S. Kabbara, et al., Diversity and evolution of sensor histidine kinases in Eukaryotes. Genome Biol. Evol. 11, 86–108 (2019).

60. H. M. Berman, et al., The cAMP binding domain: an ancient signaling module. Proceedings of the National Academy of Sciences 102, 45–50 (2005).

61. J. H. Andersen, et al., Long-term temporal and spatial trends in eutrophication status of the Baltic Sea. Biol. Rev. Camb. Philos. Soc. 92, 135–149 (2017).

62. W. R. Roberts, E. C. Ruck, K. M. Downey, E. Pinseel, A. J. Alverson, Resolving marine–freshwater transitions by diatoms through a fog of gene tree discordance. *Systematic Biology*, syad038 (2023).

63. A. J. Cortés, F. López-Hernández, D. Osorio-Rodriguez, Predicting thermal adaptation by looking into populations’ genomic past. Front. Genet. 11, 564515 (2020).

64. E. Pinseel, et al., Extinction of austral diatoms in response to large-scale climate dynamics in Antarctica. Science Advances 7, eabh3233 (2021).

65. S. Trubovitz, D. Lazarus, J. Renaudie, P. J. Noble, Marine plankton show threshold extinction response to Neogene climate change. Nat. Commun. 11, 5069 (2020).

66. H. Li, Aligning sequence reads, clone sequences and assembly contigs with BWA-MEM. arXiv [q-bio.GN] (2013).

67. R. Kofler, R. V. Pandey, C. Schlötterer, PoPoolation2: identifying differentiation between populations using sequencing of pooled DNA samples (Pool-Seq). Bioinformatics 27, 3435–3436 (2011).

68. M. Kapun, et al., Genomic analysis of European *Drosophila melanogaster* populations reveals longitudinal structure, continent-wide selection, and previously unknown DNA viruses. Mol. Biol. Evol. 37, 2661–2678 (2020).

69. M. Gautier, R. Vitalis, L. Flori, A. Estoup, f-Statistics estimation and admixture graph construction with Pool-Seq or allele count data using the R package poolfstat. Mol. Ecol. Resour. 22, 1394–1416 (2022).

70. R. Kofler, et al., PoPoolation: a toolbox for population genetic analysis of next generation sequencing data from pooled individuals. PLoS One 6, e15925 (2011).

71. P. Cingolani, et al., A program for annotating and predicting the effects of single nucleotide polymorphisms, SnpEff: SNPs in the genome of *Drosophila melanogaster* strain w1118; iso-2; iso-3. Fly 6, 80–92 (2012).

72. A. Alexa, J. Rahnenführer, Gene set enrichment analysis with topGO. Bioconductor Improv 27 (2009).

73. R. R. L. Guillard, C. J. Lorenzen, Yellow-green algae with chlorophyllide c 1, 2. J. Phycol. 8, 10–14 (1972).

74. R. R. L. Guillard, P. E. Hargraves, *Stichochrysis immobilis* is a diatom, not a chrysophyte. Phycologia 32, 234–236 (1993).

75. W. H. C. F. Kooistra, et al., Global diversity and biogeography of *Skeletonema* species (Bacillariophyta). Protist 159, 177–193 (2008).

76. J. Chuan, A. Zhou, L. R. Hale, M. He, X. Li, Atria: an ultra-fast and accurate trimmer for adapter and quality trimming. Gigabyte, 10.46471/gigabyte.31 (2021).

77. H. Li, et al., The Sequence Alignment/Map format and SAMtools. Bioinformatics 25, 2078–2079 (2009).

78. G. A. Van der Auwera, B. D. O’Connor, Genomics in the Cloud: Using Docker, GATK, and WDL in Terra (1st Edition) (O’Reilly Media, 2020).

79. O. Tange, “GNU Parallel 2018, Mar 2019, ISBN 9781387509881, doi: 10.5281/zenodo.1146014” (2018).

80. S. J. Micheletti, S. R. Narum, Utility of pooled sequencing for association mapping in nonmodel organisms. Mol. Ecol. Resour. 18, 825–837 (2018).

81. P. Vries, K. Döös, Calculating lagrangian trajectories using time-dependent velocity fields. J. Atmos. Ocean. Technol. 18, 1092–1101 (2001).

82. R. Hordoir, et al., Nemo-Nordic 1.0: a NEMO-based ocean model for the Baltic and North seas – research and operational applications. Geosci. Model Dev. 12, 363–386 (2019).

83. S. M. Uppala, et al., The ERA-40 re-analysis. Quart. J. Roy. Meteor. Soc. 131, 2961–3012 (2005).

84. S. Levitus, T. P. Boyer, World ocean atlas, vol 5, salinity: NOAA atlas (1994).

85. A. Godhe, et al., Seascape analysis reveals regional gene flow patterns among populations of a marine planktonic diatom. Proc. Biol. Sci. 280, 20131599 (2013).

86. J. W. Hurrell, C. Deser, North Atlantic climate variability: The role of the North Atlantic Oscillation. J. Mar. Syst. 78, 28–41 (2009).

87. P. R. Jonsson, P.-O. Moksnes, H. Corell, E. Bonsdorff, M. Nilsson Jacobi, Ecological coherence of marine protected areas: new tools applied to the Baltic Sea network. Aquat. Conserv. 30, 743–760 (2020).

88. J. van Etten, R Package gdistance: Distances and Routes on Geographical Grids. J. Stat. Softw. 76, 1–21 (2017).

89. S. Dray, A.-B. Dufour, The ade4 package: implementing the duality diagram for ecologists. J. Stat. Softw. 22, 1–20 (2007).

90. E. J. Pebesma, Multivariable geostatistics in S: the gstat package. Comput. Geosci. 30, 683–691 (2004).

91. J. Oksanen, et al., vegan: Community Ecology Package. R package version 2.6-2. https://CRAN.R-project.org/package=vegan (2022).

92. S. Nadeau, P. G. Meirmans, S. N. Aitken, K. Ritland, N. Isabel, The challenge of separating signatures of local adaptation from those of isolation by distance and colonization history: The case of two white pines. Ecol. Evol. 6, 8649–8664 (2016).

93. B. Leroy, C. N. Meynard, C. Bellard, F. Courchamp, virtualspecies, an R package to generate virtual species distributions. Ecography 39, 599–607 (2016).

94. Y. A. Alshwairikh, et al., Influence of environmental conditions at spawning sites and migration routes on adaptive variation and population connectivity in Chinook salmon. Ecol. Evol. 11, 16890–16908 (2021).

95. S. F. Altschul, W. Gish, W. Miller, E. W. Myers, D. J. Lipman, Basic local alignment search tool. J. Mol. Biol. 215, 403–410 (1990).

96. P. Jones, et al., InterProScan 5: genome-scale protein function classification. Bioinformatics 30, 1236–1240 (2014).

97. T. Aramaki, et al., KofamKOALA: KEGG Ortholog assignment based on profile HMM and adaptive score threshold. Bioinformatics 36, 2251–2252 (2020).

